# TGM2-mediated histone serotonylation is an epigenetic cardioprotective mechanism in HFpEF

**DOI:** 10.64898/2026.06.25.734596

**Authors:** Ryo Ogawara, Tomofumi Misaka, Yoshinori Suzuki, Satoshi Okochi, Shohei Ichimura, Shunsuke Miura, Tetsuro Yokokawa, Shu Taira, Satoshi Waguri, Masayoshi Oikawa, Akiomi Yoshihisa, Takafumi Ishida, Yasuchika Takeishi

**Author notes:** Correspondence: Tomofumi Misaka, MD, PhD, Department of Cardiovascular Medicine, Fukushima Medical University 1, Hikarigaoka, Fukushima, 960-1295, Japan.

## Abstract

Heart failure with preserved ejection fraction (HFpEF) is a heterogeneous syndrome with incompletely understood molecular mechanisms. Histone serotonylation is a recently identified epigenetic modification in which serotonin is covalently conjugated to glutamine 5 of histone H3 in H3K4me3-marked nucleosomes. Here, we investigated the role of transglutaminase 2 (TGM2)-mediated histone serotonylation in HFpEF. In a mouse model of HFpEF induced by salty drinking water, unilateral nephrectomy and aldosterone infusion (SAUNA), cardiac H3K4me3Q5ser and nuclear TGM2 levels were increased. Cardiomyocyte-specific TGM2-deficient mice developed aggravated HFpEF phenotypes after SAUNA exposure, including worsened diastolic dysfunction, reduced exercise capacity, pulmonary congestion and delayed cardiomyocyte relaxation. CUT&RUN sequencing identified H3K4me3Q5ser-enriched regions predominantly around transcription start sites after SAUNA exposure, with notable enrichment at genes associated with G2/M checkpoint-related stress-response signaling. RNA sequencing further showed that activation of this pathway was impaired in SAUNA-exposed TGM2-deficient hearts. In cardiac myocytes, calcium-binding sites and nuclear localization of TGM2 support checkpoint-related stress-response gene activation in cardiac myocytes. Pharmacological WEE1 inhibition, which activates downstream CDK1-associated checkpoint signaling, partially rescued the aggravated HFpEF phenotype in TGM2-deficient mice. Finally, in patients with HFpEF, lower circulating serotonin levels were associated with adverse cardiac outcomes, and cardiomyocyte H3K4me3Q5ser levels correlated with serum serotonin concentrations. These findings suggest that cardiomyocyte TGM2-mediated histone serotonylation represents a stress-adaptive, cardioprotective epigenetic mechanism in HFpEF.

## Introduction

Heart failure with preserved ejection fraction (HFpEF) is a heterogeneous clinical syndrome that accounts for approximately 50% of all heart failure cases and its prevalence is increasing.^1^ Impaired relaxation kinetics and increased passive myocardial stiffness are hallmarks of HFpEF and are considered major contributors to its pathophysiology.^2^ Recently, sarcomere-level contractile and relaxation abnormalities have been reported to be associated with obesity-related HFpEF phenotypes.^3^ Although the development of HFpEF is thought to be largely influenced by acquired factors,^4^ the underlying molecular mechanisms remain to be elucidated.

Epigenetic mechanisms play a key role in various types of disease development by linking environmental and pathological stimuli to changes in gene expression without altering the underlying DNA sequence.^5^ Therapeutic strategies targeting epigenetic regulation, including histone modifications, have been extensively explored in oncology.^6^ In cardiovascular disease, particularly heart failure with reduced ejection fraction, differential DNA methylation and certain histone modifications, such as histone acetylation and deacetylation, have been reported,^7^ and histone deacetylase 4 has been shown to exert a protective role against the development of heart failure.^8^ While targeting epigenetic regulation may represent a promising therapeutic approach for heart failure, the contribution of histone modifications to HFpEF pathogenesis has remained largely unexplored and such therapies have not yet been established.

Histone serotonylation has been recently identified as a novel histone post-translational modification.^9^ Serotonylation of glutamine occurs at position 5 (Q5ser) on histone H3 in the presence of tri-methylated lysine 4 (H3K4me3)-marked nucleosomes, resulting in the combinatorial mark H3K4me3Q5ser. Histone serotonylation is mediated by the transglutaminase activity of transglutaminase 2 (TGM2).^9^ Although the functional role of H3K4me3Q5ser has been characterized in neurons, where it regulates stress-induced transcriptional plasticity in rodent models of major depressive disorder,^9,10^ its role in the cardiovascular system remains largely unknown. Furthermore, although intracellular signaling mediated by membrane receptors, particularly the serotonin 2B receptor, has been reported in the context of serotonin as a neurotransmitter,^11,12^ the overall role of serotonin in the heart remains incompletely understood; notably, its epigenetic functions within the nucleus of cardiac cells have largely been overlooked.

Here, we demonstrate the role of histone serotonylation in the pathogenesis of HFpEF, and propose that targeting histone serotonylation may be a potential approach to prevent HFpEF development.

## Results

### Alteration of cardiac histone serotonylation levels during the development of HFpEF

To clarify the role of histone serotonylation in the development of HFpEF, we established a salt water, unilateral nephrectomy, and aldosterone infusion (SAUNA) model in C57BL/6 mice, which integrates multiple cardiometabolic and hemodynamic stressors and closely mimics the complex pathophysiology of human HFpEF (Figure 1A).^13,14,15^ Four weeks after SAUNA, SAUNA-exposed mice exhibited increased blood pressure, elevated serum creatinine and urinary albumin levels, and impaired diastolic function, as evidenced by increased E/A and E/e′ ratios and left atrial volume whereas fractional shortening was preserved (Supplementary Figure 1A-1E). In addition, these mice showed reduced running distance and increased left ventricular weight and lung weight, as well as mRNA expressions of *Nppa*, *Myh7*, and *Col3a1* in SAUNA hearts (Supplementary Figure 1F-1H). Because histone serotonylation is catalyzed by TGM2,^9^ we next examined cardiac H3K4me3Q5ser levels and TGM2 localization in SAUNA hearts. Cardiac histone serotonylation levels, as indicated by H3K4me3Q5ser, were significantly elevated compared to those in the sham group (Figure 1B and 1C). Nuclear TGM2 levels in SAUNA hearts were significantly elevated compared to sham hearts whereas cytosolic TGM2 remained unchanged (Figure 1B and 1C). Stimulation with aldosterone elevated H3K4me3Q5ser levels in neonatal rat cardiomyocytes (NRCM) and H9c2 cardiac myocytes (Figure 1D and Supplementary Figure 2A). In contrast, treatment of the TGM2 inhibitor, LDN27219, suppressed the aldosterone-induced increase in H3K4me3Q5ser levels in NRCM and H9c2 myocytes (Figure 1E and Supplementary Figure 2B). Additionally, serotonin stimulation increased H3K4me3Q5ser levels (Supplementary Figure 2C). We analyzed publicly available single-nucleus sequencing datasets from the human heart.^16^ We found that *TGM2* expression was enriched in cardiomyocytes (Figure 1F and 1G). Moreover, *TGM2* transcript levels were decreased in patients with heart failure due to hypertrophic cardiomyopathy compared to non-heart failure controls (Figure 1H). These findings indicate that cardiac TGM2 is dynamically regulated at multiple levels, including subcellular localization and gene expression, and suggest that histone serotonylation may contribute to the epigenetic reprogramming underlying HFpEF pathogenesis.

**Figure 1.**
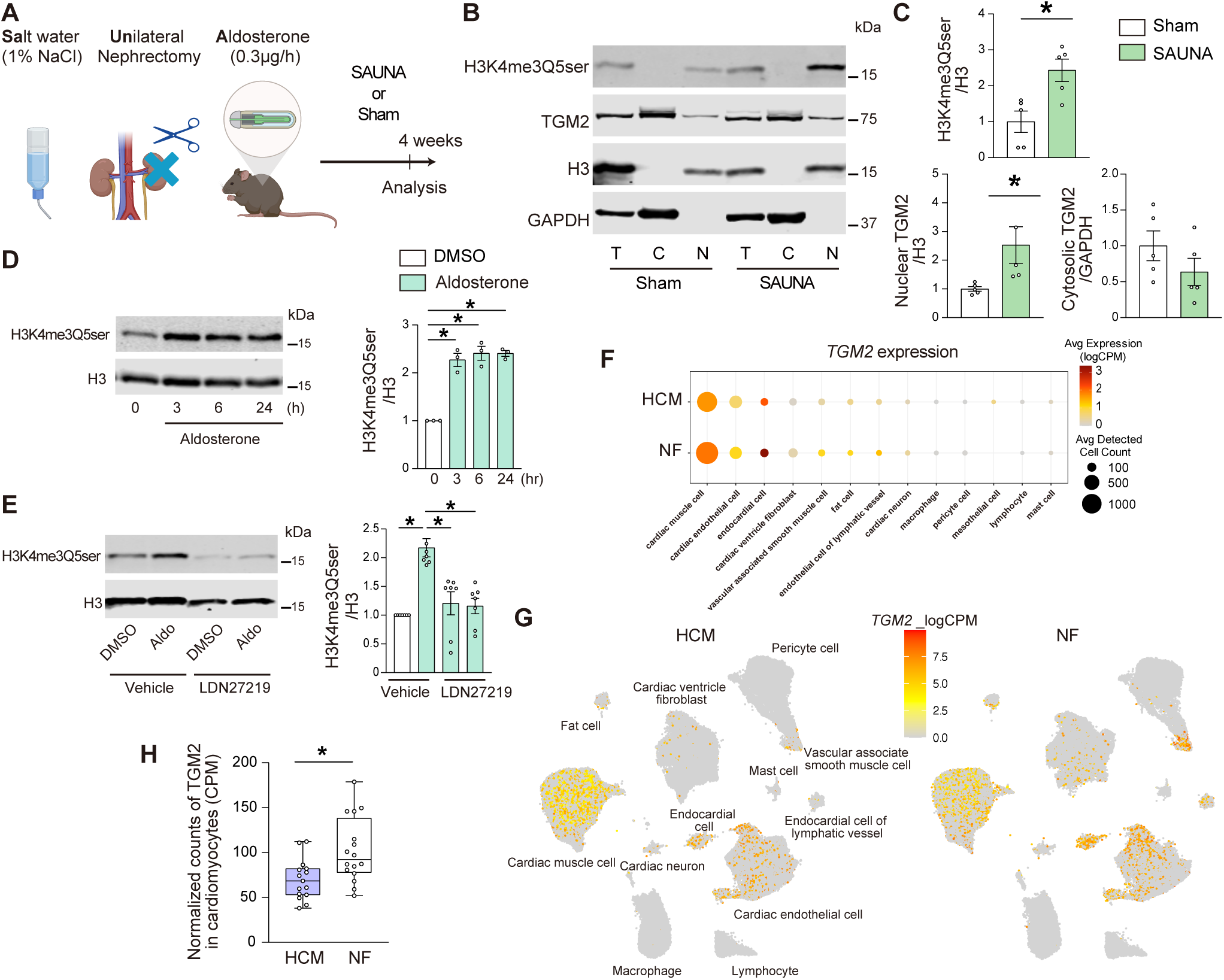
Alteration of cardiac histone serotonylation levels during the development of HFpEF. **A**, Schematic illustration of the SAUNA protocol, consisting of 1% saline drinking water, unilateral nephrectomy, and continuous aldosterone infusion (0.30 µg/h). C57BL/6 wild-type (WT) mice underwent either SAUNA or sham and analyzed at 4 weeks after initiation. **B**, Expression levels of H3K4me3Q5ser and TGM2 in the nuclear and cytosolic fractions of the hearts following SAUNA. H3 and GAPDH were used for nuclear and cytosolic loading controls, respectively. T represents total protein; C, cytosolic fraction; N, nuclear fraction. **C**, Quantitative densitometric analyses of the immunoblots from (B) are presented in the graphs (n = 5). **D**, H3K4me3Q5ser levels in cardiac cells. Neonatal rat cardiomyocytes (NRCMs) stimulated with aldosterone (100 nM) for the indicated time points (n=3). **E**, Effects of the TGM2 inhibitor LDN27219 on H3K4me3Q5ser in aldosterone-treated NRCM (n=7). **F**, Publicly available human cardiac single-nucleus RNA-sequencing analysis of *TGM2* expression. Dot plot showing *TGM2* expression across cardiac cell types in heart failure with hypertrophic cardiomyopathy (HCM) and non-failing controls (NF). Dot color indicates average expression (logCPM), and dot size indicates the number of cells in which *TGM2* transcripts were detected. **G**, UMAP plots showing the distribution of *TGM2* expression in HCM and NF hearts. **H**, Comparison of *TGM2* levels, shown as logCPM, between HCM and NF hearts. Data are presented as mean ± SEM or median with interquartile range, as appropriate. P < 0.05 by two-sided unpaired t test, one-way ANOVA with Tukey post hoc test, or Mann–Whitney U test.

### Deficiency of cardiac TGM2 exacerbates SAUNA-induced HFpEF phenotypes

To elucidate the role of histone serotonylation in the development of HFpEF in vivo, we generated cardiomyocyte-specific TGM2 knockout (TGM2-CKO) mice. Homozygous *Tgm2* flox mice (*Tgm2*^flox/flox^) were crossed with αMHC-Cre transgenic mice (*Tgm2*^flox/flox^; αMHC-Cre^+^). *Tgm2*^flox/flox;^αMHC-Cre^−^ littermates were used as wild-type (WT) controls. TGM2-CKO mice were born at the expected Mendelian ratio (53 and 51 mice, respectively), grew normally to adulthood, and were fertile. The protein level of TGM2 significantly decreased in TGM2-CKO hearts by 80% compared to WT littermates (Figure 2A). H3K4me3Q5ser levels were significantly decreased in TGM2-CKO hearts by 78% (Figure 2B).

**Figure 2.**
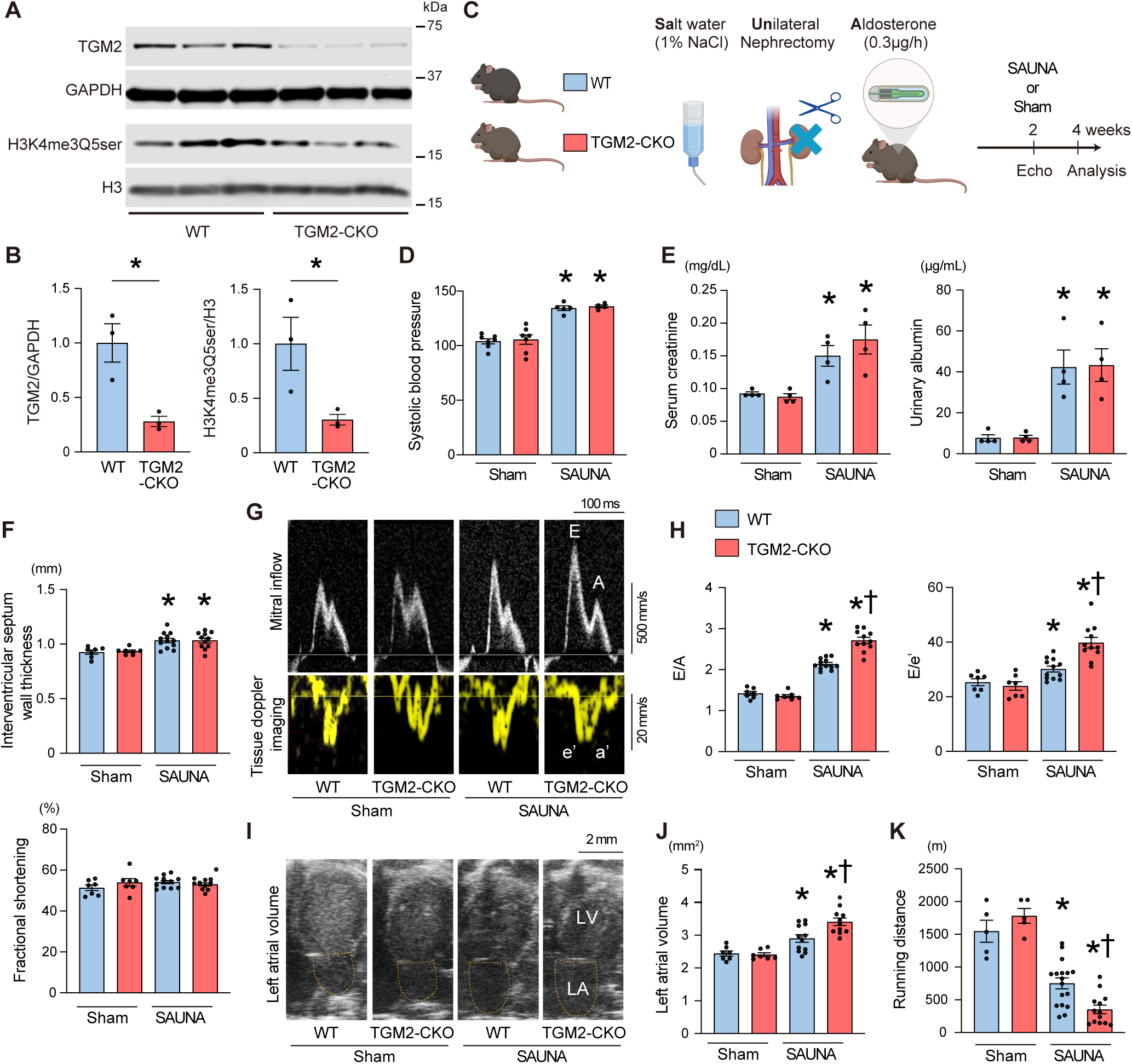
Deficiency of cardiac TGM2 exacerbates SAUNA-induced HFpEF phenotypes. **A**, Protein expression levels of TGM2 and H3K4me3Q5ser in the hearts of cardiomyocyte-specific TGM2 knockout (TGM2-CKO) mice and WT littermates. **B**, Densitometric analysis of immunoblots from (A). **C**, Schematic illustration of the experimental protocol. TGM2-CKO mice and WT mice were subjected to SAUNA or sham protocol and analyzed at 4 weeks after initiation. **D**, Systolic blood pressure. **E**, Serum creatinine and urinary albumin levels. **F**, Echocardiographic parameters. **G**, Representative echocardiographic images of pulsed-wave Doppler and tissue Doppler imaging. Peak early (E) and late (A) transmitral inflow velocities and early (e’) and late (a) diastolic mitral annular velocity are shown. **H**, E/A and E/e’ ratios. **I**, Representative echocardiographic images of left atrial volume. LA and LV indicate left atrium and left ventricle, respectively. **J**, Quantification of the left atrial volume. **K**, Running distance determined by treadmill exercise testing. All data are presented as mean□±□SEM. *P□<□0.05 versus the corresponding sham-exposed group and ^†^P□<□0.05 versus the corresponding WT mice by the one-way ANOVA with Tukey post hoc analysis.

To examine the role of histone serotonylation and TGM2 under HFpEF conditions, TGM2-CKO mice were subjected to the SAUNA protocol (Figure 2C). Four weeks after SAUNA, systolic, mean, and diastolic blood pressure were elevated, along with increases in serum creatinine, blood urea nitrogen, urinary albumin levels, but these parameters did not differ significantly between WT and TGM2-CKO mice (Figure 2D and 2E, Supplementary Figure 3A-3D). In echocardiography, end-diastolic interventricular septum wall thickness was increased in response to SAUNA, but this did not differ significantly between two groups, and fractional shortening was preserved (Figure 2F). Regarding diastolic function, SAUNA increased E/A, E/e’ ratios, and left atrial volume in both WT and TGM2-CKO mice; notably, TGM2-CKO mice showed significantly greater elevations in these parameters compared to WT mice (Figure 2G-2J). We observed that TGM2-CKO mice exhibited increased E/A and E/e’ ratios at the 2-week time point in response to SAUNA (Supplementary Figure 4). Importantly, treadmill exercise testing demonstrated a significant reduction in running distance in both WT mice and TGM2-CKO mice after SAUNA, but TGM2-CKO mice showed a further reduction in running distance (Figure 2K).

**Figure 3.**
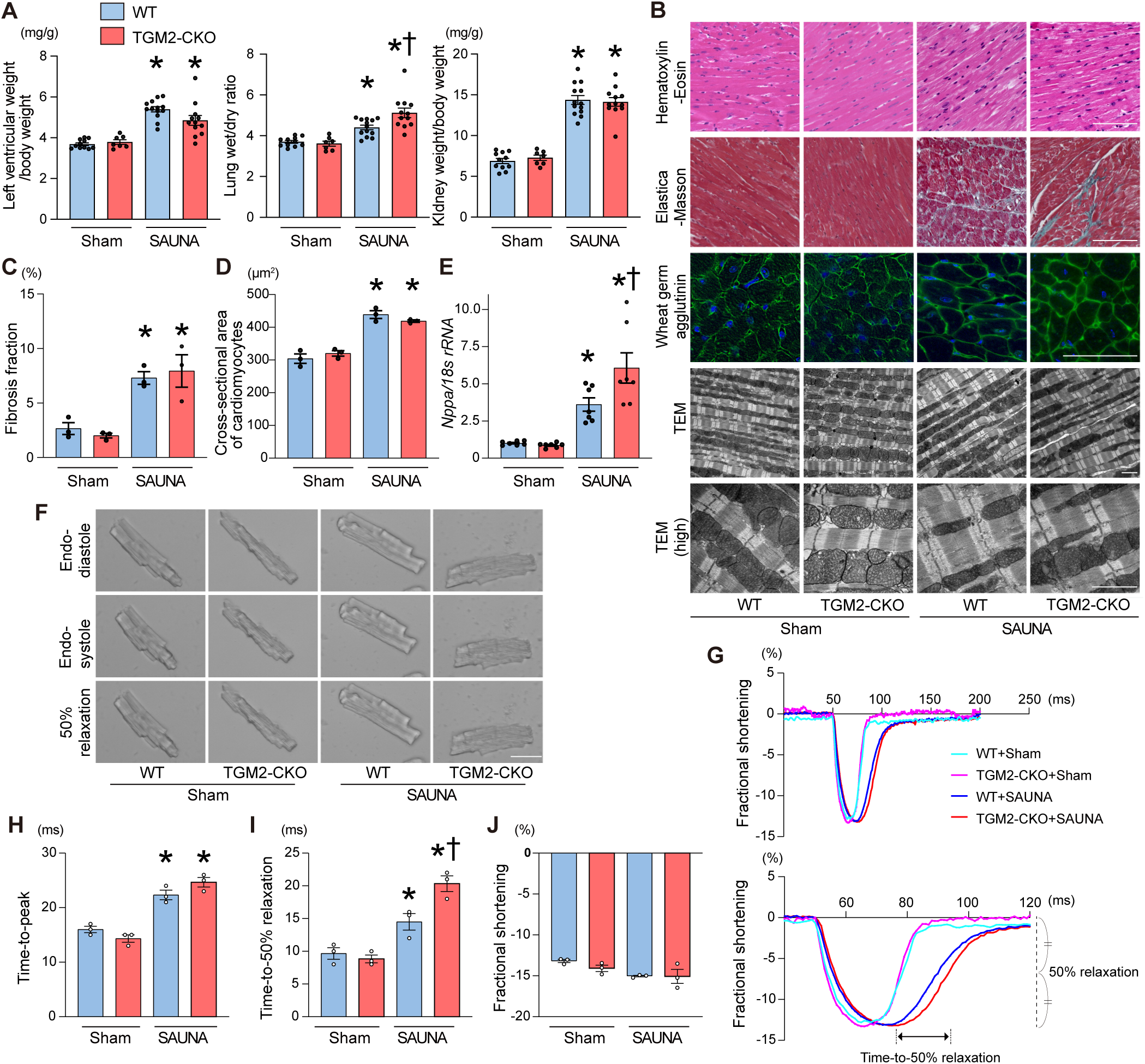
Knockout of TGM2 leads to delayed relaxation kinetics in isolated cardiomyocytes. **A**, Physiological parameters of sham or SAUNA-exposed WT and TGM2-CKO mice. **B**, Representative images of Hematoxylin–Eosin staining, Elastica–Masson staining, wheat germ agglutinin (WGA; green) with DAPI (blue) staining, and transmission electron microscopy (TEM) of left ventricular sections. Scale bars: 100 μm for histological and fluorescence images; 2 μm for TEM images. **C**, Quantitative assessment of myocardial fibrosis based on EM staining (n = 3 per group). **D**, Quantification of cardiomyocyte cross-sectional area measured from WGA-stained sections (n = 3 per group). **E**, Relative mRNA expression levels of *Nppa* of the hearts. Gene expression was normalized to *18s rRNA*, and values are presented relative to the sham group, which was set to 1 (n = 7). **F**, Representative images of isolated cardiomyocytes from WT and TGM2-CKO mice after sham or SAUNA at end-diastole, end-systole, and 50% relaxation during 1-Hz electrical pacing. **G**, Representative sarcomere shortening traces. Lower traces show the same recordings displayed on an expanded time scale to visualize relaxation kinetics. Time-to-50% relaxation in SAUNA-exposed TGM2-CKO cardiomyocyte is indicated. **H-J**, Quantification of sarcomere shortening kinetics in isolated cardiomyocytes from each group. **H**, Time to peak shortening. **I**, Time to 50% relaxation. **J**, Fractional shortening (n = 3 per group). *P□<□0.05 versus the corresponding sham-exposed group and ^†^P□<□0.05 versus the corresponding WT mice by the one-way ANOVA with Tukey post hoc analysis.

### Knockout of TGM2 leads to delayed relaxation kinetics at a single cardiomyocyte level

After tissue harvesting, lung weight and lung wet-to-dry weight ratio were elevated following SAUNA induction; however, these indices of pulmonary congestion were significantly greater in SAUNA-exposed TGM2-CKO mice than in SAUNA-exposed WT mice (Figure 3A and Supplementary Figure 3E). Whole heart weight and left ventricular weight, as well as kidney weight were increased after SAUNA exposure but did not differ between the two groups. Histological analysis showed SAUNA increased interstitial fibrosis as well as cross-sectional area of cardiomyocytes, but no significant differences were observed between SAUNA-exposed WT and TGM2-CKO mice (Figure 3B-3D). Electron microscopic analysis revealed that sarcomeric structure was preserved, with no apparent disruption of myofibrillar organization across groups (Figure 3B). mRNA expression of *Nppa* was significantly higher in SAUNA-exposed TGM2-CKO hearts than in SAUNA-exposed WT hearts (Figure 3E). Mass spectrometry imaging revealed a trend toward reduced 5-hydroxytryptamine (5-HT) signal intensity in the left ventricle of SAUNA-exposed TGM2-CKO mice (Supplementary Figure 5). Taken together, these findings indicate that the absence of cardiac TGM2 exacerbates diastolic cardiac function, reduces exercise capacity, and increases pulmonary congestion under SAUNA conditions, without altering hypertrophic response or myocardial fibrosis.

Then, we investigated relaxation kinetics at the single cardiomyocyte level. Cardiomyocytes were isolated and cultured from WT and TGM2-CKO mice following SAUNA or sham exposure, and cell shortening and relaxation kinetics were assessed in response to 1-Hz electrical pacing (Supplementary Video).^17,18^ SAUNA exposure prolonged time-to-peak and time-to-50% relaxation in cardiomyocytes compared with sham exposure (Figure 3F-3I). Notably, the time-to-50% relaxation was significantly prolonged in SAUNA-exposed TGM2-CKO cardiomyocytes compared to SAUNA-exposed WT cardiomyocytes, while time-to-peak shortening was not different between them. Fractional shortening did not differ among the four groups (Figure 3J). These findings indicate that the absence of cardiomyocyte TGM2 exacerbates SAUNA-induced impairment of relaxation capacity at the single-cell level.

### Genome-wide effects of histone serotonylation in HFpEF

To investigate the genome-wide distribution and regulatory effects of histone serotonylation, we performed CUT&RUN sequencing using an anti-H3K4me3Q5ser antibody in hearts from sham- and SAUNA-exposed WT mice (Figure 4A). Given that cardiomyocyte-specific TGM2 deletion resulted in a marked reduction of H3K4me3Q5ser, CUT&RUN with anti-H3K4me3Q5ser antibody did not yield sufficient enrichment of immunoprecipitated DNA in TGM2-CKO hearts. CUT&RUN analysis identified reproducible H3K4me3Q5ser-enriched regions across biological replicates,^19^ indicating that this histone serotonylation mark is detectable in cardiac chromatin. The numbers of H3K4me3Q5ser consensus peaks were 5,093 and 11,718 in sham and SAUNA hearts, respectively (Figure 4B). Metagene and heatmap analysis further demonstrated prominent H3K4me3Q5ser enrichment around transcription start sites, supporting the notion that H3K4me3Q5ser is associated with transcriptionally active chromatin regions (Figure 4C). Next, to gain insight into the molecular features of genes associated with SAUNA-induced H3K4me3Q5ser-enriched regions, we performed gene set enrichment analysis (GSEA) using peak-associated genes. This analysis identified significant enrichment of several HALLMARK pathways, including IL2/STAT5 signaling, G2/M checkpoint, and p53 pathway (Figure 4D).

**Figure 4.**
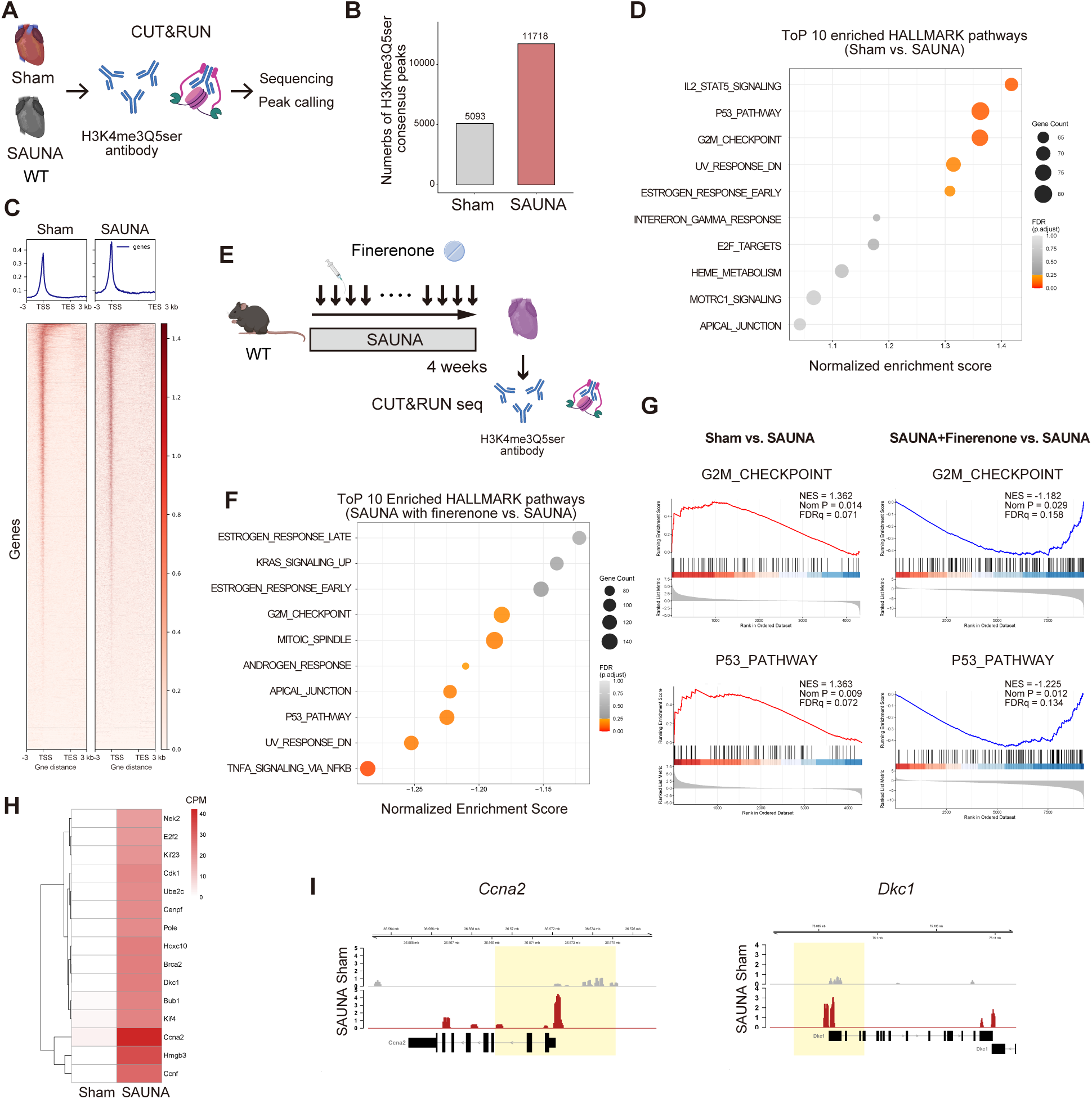
Genome-wide profiling of H3K4me3Q5ser in HFpEF hearts. **A**, Schematic experimental protocol of CUT&RUN sequencing using an H3K4me3Q5ser antibody in the hearts from sham and SAUNA-exposed mice. **B**, Number of H3K4me3Q5ser consensus peaks identified in sham and SAUNA groups. **C**, Heatmaps and metagene profiles showing H3K4me3Q5ser signal intensity around gene bodies in sham and SAUNA hearts. Distribution of H3K4me3Q5ser signals across gene bodies, from 3 kb upstream of transcription start sites (TSS) to 3 kb downstream of transcription end sites (TES). **D**, Top enriched HALLMARK pathways associated with SAUNA-induced H3K4me3Q5ser changes compared with sham. Pathways are ranked by normalized enrichment score. Dot size indicates gene count, and color indicates adjusted P value. **E**, Experimental scheme for finerenone treatment and H3K4me3Q5ser CUT&RUN sequencing. Finerenone was administered orally once daily at a dose of 10 mg/kg body weight for 4 weeks in SAUNA-exposed WT mice. **F**, Top HALLMARK pathways altered by finerenone treatment in SAUNA mice compared with untreated SAUNA mice. Pathways are ranked by NES. Dot size indicates gene count, and color indicates adjusted P value. **G**, Gene set enrichment analysis plots showing enrichment of HALLMARK_G2M_CHECKPOINT and HALLMARK_P53_PATHWAY in SAUNA versus sham and in finerenone-treated SAUNA versus untreated SAUNA hearts. **H**, Heatmap showing H3K4me3Q5ser-associated G2/M checkpoint genes differentially enriched between sham and SAUNA hearts. **I**, Representative genome browser tracks showing H3K4me3Q5ser CUT&RUN signal at selected G2/M checkpoint-associated loci, including *Ccna2* and *Dkc1* in sham and SAUNA hearts. Yellow shading indicates TSS ± 3kb.

To determine whether the epigenomic alterations observed in SAUNA-exposed hearts were therapeutically reversible and to identify the pathways targeted by HFpEF-associated H3K4me3Q5ser-enriched loci, we treated SAUNA-exposed mice with finerenone, a nonsteroidal mineralocorticoid receptor antagonist,^20^ for 4 weeks (Figure 4E). Finerenone treatment ameliorated SAUNA-induced diastolic dysfunction, improved exercise capacity, and reduced cardiac hypertrophy and pulmonary congestion (Supplementary Figure 6). We then compared H3K4me3Q5ser enrichment and HFpEF-associated transcriptional signatures between finerenone-treated and untreated SAUNA hearts (Figure 4F). Differentially enriched H3K4me3Q5ser peaks were preferentially associated with genes involved in the G2/M checkpoint and p53 pathways, and these SAUNA-induced epigenomic alterations were attenuated by finerenone treatment (Figure 4G). In addition, heatmap analysis of the top 15 G2/M checkpoint-associated genes demonstrated a coordinated increase in H3K4me3Q5ser-associated signals in the SAUNA group compared to sham (Figure 4H and 4I). These findings indicate that SAUNA induces H3K4me3Q5ser accumulation at genomic loci linked to cellular stress responses.

### G2/M checkpoint–related gene expression is reduced in TGM2 knockout hearts after SAUNA exposure

To identify functionally relevant pathways associated with the TGM2-knockout phenotype in HFpEF, we analyzed RNA-sequence data from cardiac tissues to characterize transcriptional changes linked to TGM2-mediated histone serotonylation. SAUNA exposure induced enrichment of G2/M checkpoint, E2F targets, mitotic spindle, inflammatory signaling, and interferon response pathways in WT hearts (Supplementary Figure 8A and 8B). Of note, GSEA comparing SAUNA-exposed WT and TGM2-CKO hearts revealed that the HALLMARK_G2M_CHECKPOINT pathway was one of the most significantly downregulated pathways in TGM2-CKO hearts after SAUNA exposure (Figure 5A and Supplementary Figure 8C). E2F_TARGETS and MITOTIC_SPINDLE were also identified. Heatmap analysis demonstrated that G2/M checkpoint–related genes were broadly upregulated in SAUNA-exposed WT hearts compared with sham controls, whereas this response was attenuated in SAUNA-exposed TGM2-CKO hearts (Figure 5B). Consistently, scatter plot analysis of log□ fold changes in HALLMARK_G2M_CHECKPOINT genes showed that a subset of genes induced by SAUNA in WT hearts was reduced or insufficiently induced in TGM2-CKO hearts (Figure 5C). GSEA enrichment curves further confirmed that SAUNA exposure activated the G2/M checkpoint gene program in WT hearts, whereas this activation was markedly blunted in TGM2-CKO hearts (Figure 5D and 5E). To identify the core genes underlying this pathway-level alteration, we extracted the overlapping 48 leading-edge genes that were induced by SAUNA in WT hearts and reduced in SAUNA-exposed TGM2-CKO hearts among 63 leading-edge genes (Figure 5F). Protein-protein interaction network analysis of these overlapping genes identified *Cdk1* and *Ccna2* as central hub genes (Figure 5G). CCNA2 and CDK1 are known to form a functional kinase complex that regulates G2/M cell-cycle progression, and both molecules have been implicated in DNA damage response and checkpoint control.^21^ Given that adult cardiomyocytes are largely post-mitotic, we interpreted the enrichment of G2/M checkpoint–related genes as reflecting a stress-responsive checkpoint program rather than productive cell-cycle progression.^22,23^ These findings suggest that TGM2-mediated histone serotonylation contributes to SAUNA-induced activation of a checkpoint-associated stress program in cardiomyocytes, thereby attenuating HFpEF progression under cardiometabolic stress.

**Figure 5.**
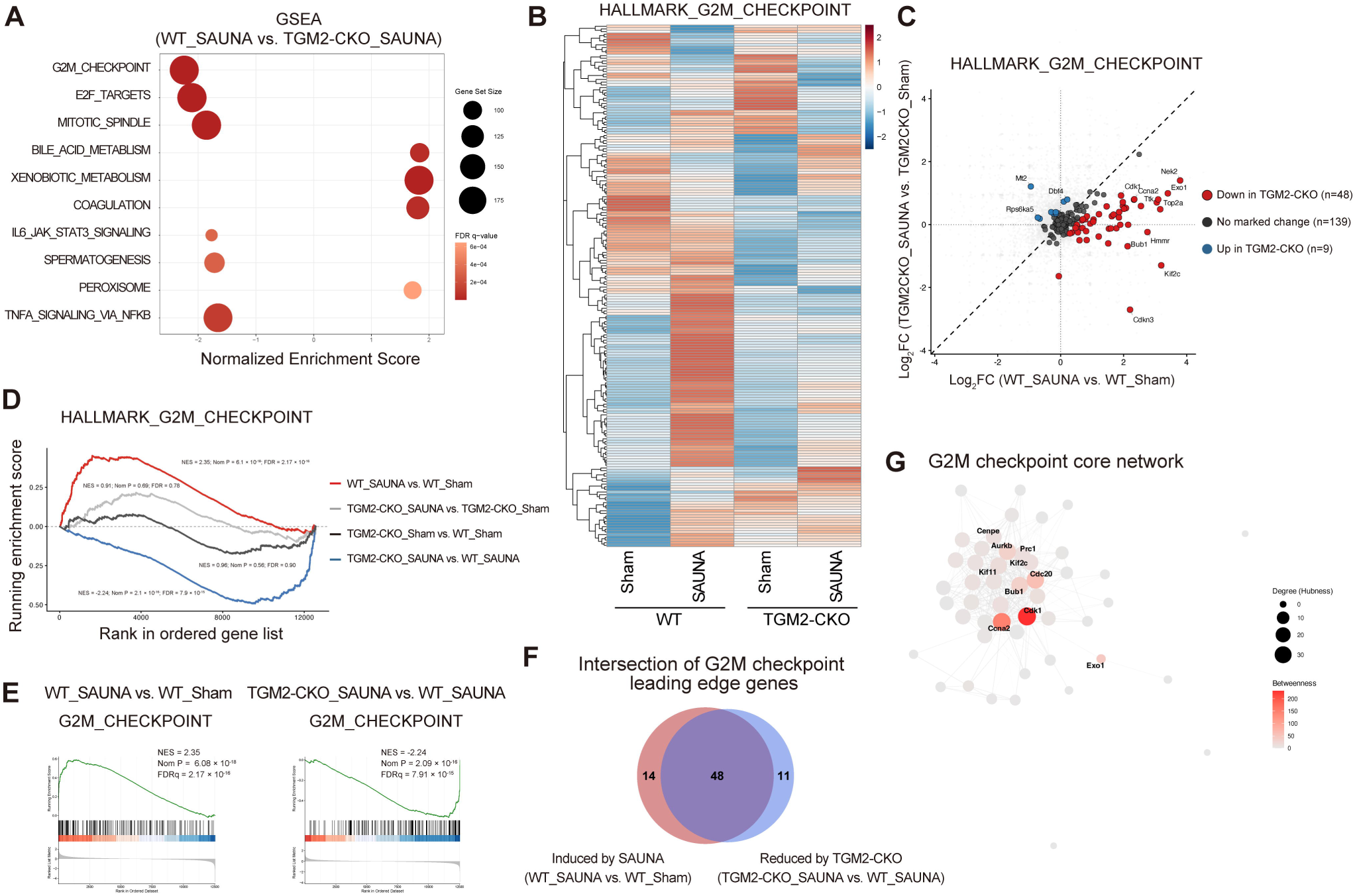
SAUNA-induced activation of G2/M checkpoint-associated transcriptional programs was attenuated in TGM2 deficiency. **A**, Gene set enrichment analysis (GSEA) of differentially expressed genes between WT_SAUNA and TGM2-CKO_SAUNA hearts by RNA-sequencing. The bubble plot shows significantly enriched Hallmark gene sets ranked by normalized enrichment score (NES) and negative NES indicates reduced enrichment in TGM2-CKO SAUNA hearts compared with WT SAUNA hearts. Bubble size indicates gene set size, and color indicates FDR q-value. **B**, Heatmap showing the expression pattern of genes included in the HALLMARK_G2M_CHECKPOINT gene set across WT and TGM2-CKO hearts under sham or SAUNA conditions. Gene expression values are scaled by row. **C**, Scatter plot comparing log□ fold changes of HALLMARK_G2M_CHECKPOINT genes in WT_SAUNA versus WT_Sham and TGM2-CKO_SAUNA versus TGM2-CKO_Sham. Genes induced by SAUNA in WT but reduced or not induced in TGM2-CKO are highlighted. **D**, GSEA enrichment curves for the HALLMARK_G2M_CHECKPOINT gene set across the indicated comparisons. SAUNA markedly enriched the G2M checkpoint pathway in WT mice, whereas this enrichment was attenuated in TGM2-CKO mice. **E**, Representative GSEA plots of the HALLMARK_G2M_CHECKPOINT gene set for WT_SAUNA versus WT_Sham and TGM2-CKO_SAUNA versus WT_SAUNA comparisons. **F**, Venn diagram showing the overlap between G2M checkpoint leading-edge genes induced by SAUNA in WT hearts and those reduced in TGM2-CKO_SAUNA compared with WT_SAUNA. **G**, Core network analysis of the overlapping G2M checkpoint leading-edge genes. Node size represents degree hubness, and node color indicates betweenness centrality. Hub genes, including *Cdk1* and *Ccna2*, are highlighted.

**Figure 6.**
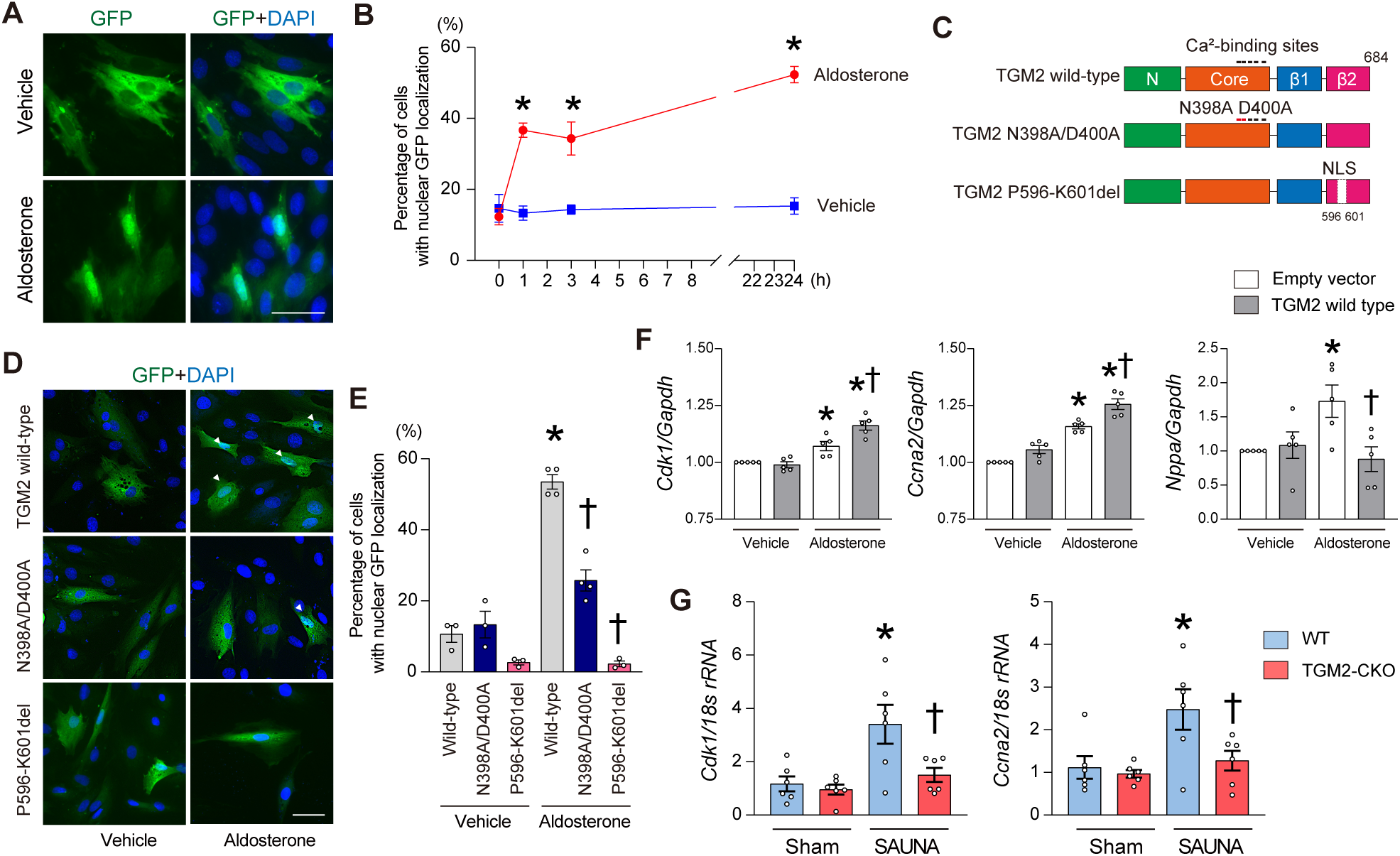
Aldosterone-induced histone serotonylation is mediated by nuclear localization of TGM2 in cardiac myocytes. **A**, Representative immunofluorescence images of GFP-tagged TGM2 in H9c2 cardiac myocytes. At 24 h after transfection with GFP-TGM2, cells were stimulated with aldosterone (100 nM) or vehicle, and fixed and stained with DAPI (blue). Scale bars, 50 μm. **B**, Quantification of the percentage of cells showing nuclear GFP localization (n=3). *P < 0.05 by two-way repeated-measures analysis of variance followed by Sidak’s multiple comparisons at the indicated time point. **C**, Schematic representation of the mouse TGM2 domain structure. N, N-terminal β-sandwich domain; Core, catalytic core domain; β1 and β2, two C-terminal β-barrel domains. Ca^2+^-binding sites and nuclear localization signal are indicated. **D**, Representative immunofluorescence images of cells expressing GFP-tagged wild-type TGM2, or the indicated TGM2 mutants, N398A/D400A and P596–K601del. At 24 h after transfection, cells were stimulated with aldosterone (100 nM) or vehicle. Scale bars, 50 μm. **E**, Quantification of percentage of cells with nuclear GFP localization (n=3-4). *P < 0.05 versus the corresponding vehicle group and ^†^P < 0.05 versus aldosterone-treated wild-type TGM2 by the one-way ANOVA with Tukey post hoc analysis. **F**, Quantification of *Cdk1*, *Ccna2*, and *Nppa* mRNA levels in H9c2 myocytes. Transfected cells with empty vector or wild-type TGM2 were stimulated with vehicle or aldosterone for 24 hours (n=5). *P < 0.05 versus the corresponding vehicle group and ^†^P < 0.05 versus aldosterone-treated empty vector by the one-way ANOVA with Tukey post hoc analysis. **G**, Quantification of *Cdk1* and *Ccna2* mRNA levels in the hearts from sham or SAUNA-exposed WT and TGM2-CKO mice. *P < 0.05 versus the corresponding sham-exposed group and ^†^P < 0.05 versus the corresponding WT mice by the one-way ANOVA with Tukey post hoc analysis.

**Figure 7.**
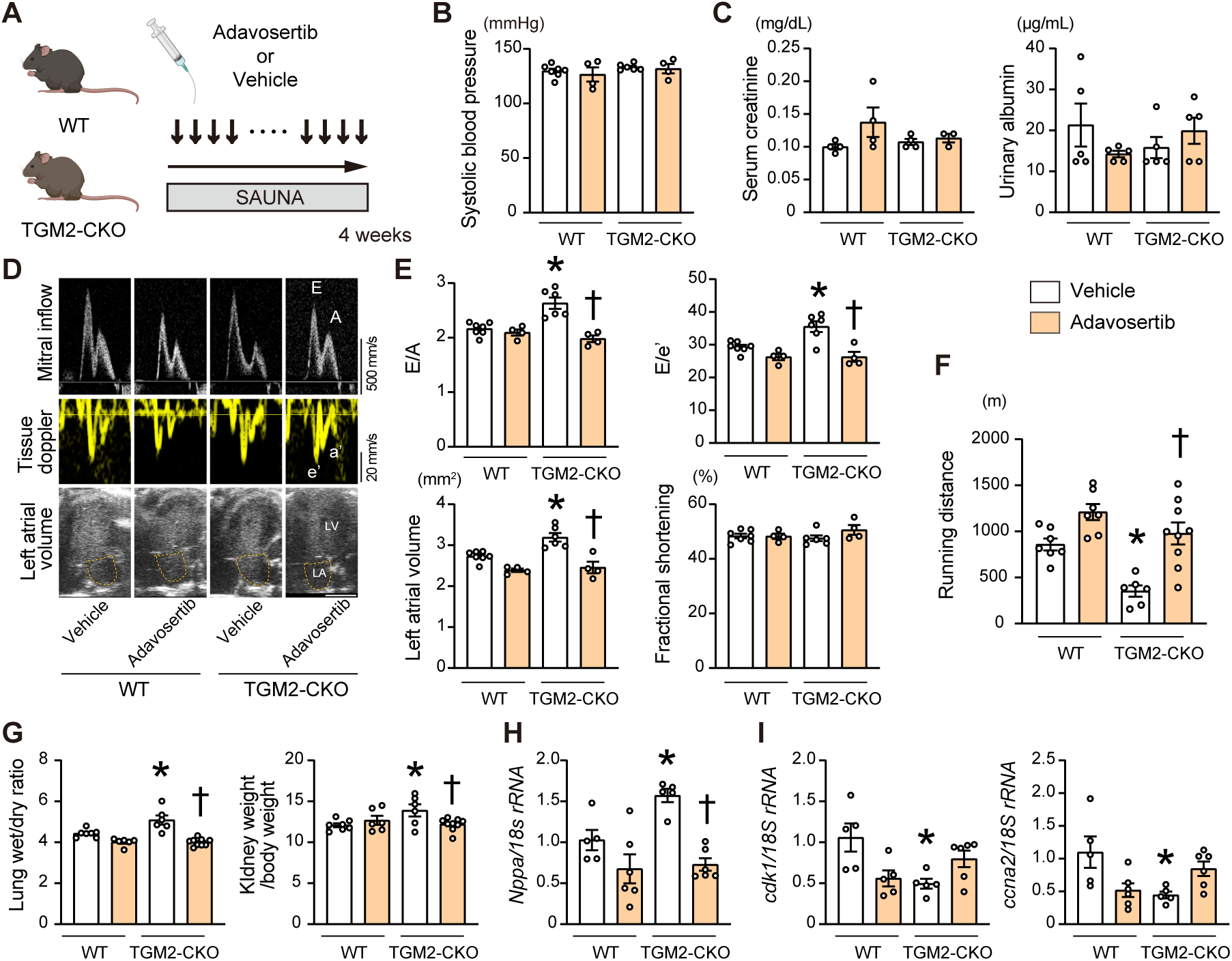
Inhibition of WEE1 improves SAUNA-induced HFpEF phenotype in TGM2-CKO mice. **A**, Schematic protocol. After initiation of the SAUNA model, the WEE1 inhibitor adavosertib (50 mg/kg/day) or vehicle was administered orally once daily for 4 weeks in WT and TGM2-CKO mice. **B**, Systolic blood pressure (n=4-7). **C**, Serum creatinine and urinary albumin levels (n=4-7). **D**, Representative echocardiographic images of pulsed- wave Doppler and tissue Doppler imaging. **E** E/A, E/e’ ratios, left atrial volume, and fractional shortening (n=4-7). **F**, Running distance determined by treadmill exercise testing (n=6-9). **G**, Physiological parameters (n=6-9). **H**, Relative mRNA expression levels of *Nppa* (n=5-6). **I**, mRNA expression levels of *cdk1* and *ccna2* (n=5-6). All data are presented as mean ± SEM. *P < 0.05 versus the vehicle-treated WT mice and ^†^P < 0.05 versus vehicle-treated TGM2-CKO mice by the one-way ANOVA with Tukey post hoc analysis.

**Figure 8.**
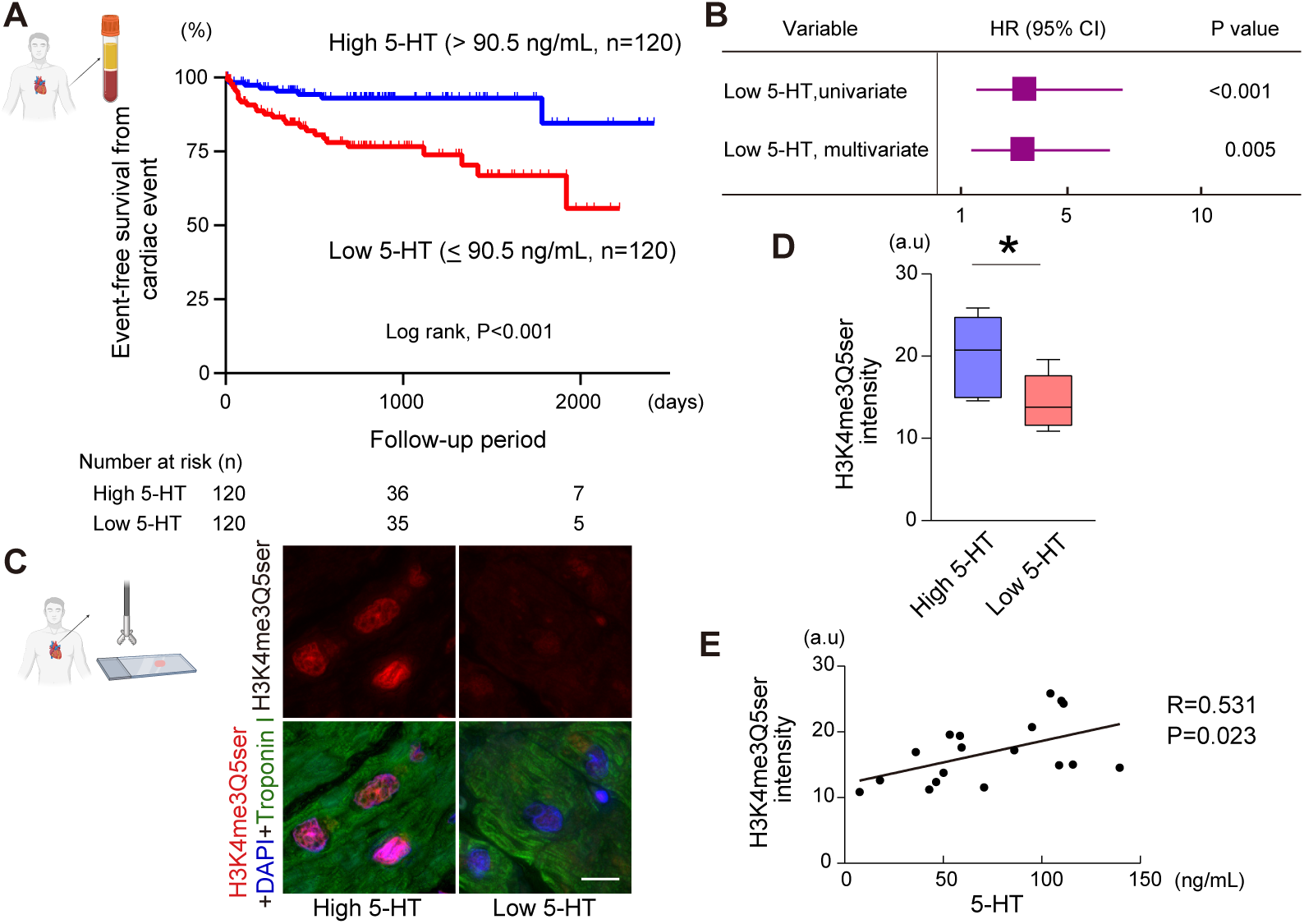
Clinical relevance of circulating serotonin and cardiomyocyte H3K4me3Q5ser in HFpEF patients. **A**, Kaplan-Meier analysis for the composite cardiovascular events for cardiac death and worsening heart failure in patients with HFpEF (n=240) stratified by serum 5-hydroxytryptamine (5-HT) levels. Patients were divided into high and low 5-HT groups based on the median value of 90.5 ng/mL. Statistical significance was assessed using the log-rank test. **B**, Forest plot showing hazard ratios from univariate and multivariate Cox proportional hazards models for the composite cardiac outcome. The multivariable model was adjusted for age, body mass index, chronic kidney disease, diuretic use, hemoglobin, and B-type natriuretic peptide. **C**, Representative immunohistochemical images of H3K4me3Q5ser (red), troponin I (green) and DAPI nuclear staining (blue) in myocardial biopsy specimens of HFpEF patients with high and low 5-HT levels. Scale bar, 20□µm. **D**, Quantification of cardiomyocyte H3K4me3Q5ser staining intensity in HFpEF patients with high (n=7) or low (n=11) serum 5-HT levels. Data are presented as median with interquartile range by Mann–Whitney U test. **E**, Correlation between serum 5-HT levels and H3K4me3Q5ser intensity in myocardial tissue (n=18). Correlation was assessed using Spearman’s rank correlation coefficient.

### Ca^2+^-binding sites and nuclear localization of TGM2 support checkpoint-related stress-response gene activation in cardiac myocytes

We then investigated how TGM2 is activated in response to HFpEF-related stress. Live-cell imaging using a GFP-tagged TGM2 expression vector demonstrated that aldosterone stimulation led to an increase in TGM2 nuclear localization (Figure 6A and 6B). Given that TGM2 contains five Ca^2+^-binding sites within catalytic core domains, we generated a mutant TGM2 vector carrying point mutations at two of Ca^2+^-binding sites, N398A and D400A (Figure 6C). This mutant, as well as deletion mutants lacking the nuclear localization signal, failed to translocate to the nucleus upon aldosterone stimulation (Figure 6D and 6E). These findings suggest that aldosterone-induced Ca^2+^-binding is essential for the nuclear localization of TGM2. Aldosterone stimulation induced expression of G2/M checkpoint–related genes, including *Cdk1*, *Ccna2*, and overexpression of TGM2 further enhanced their expressions (Figure 6F). In contrast, the cardiac remodeling marker *Nppa* was reduced by TGM2 overexpression under aldosterone stimulation. Consistently, *Cdk1* and *Ccna2* mRNA were significantly reduced in TGM2-knockout hearts after SAUNA exposure compared with WT hearts (Figure 6G). These data indicate that aldosterone promotes Ca^2+^-dependent nuclear translocation of TGM2, thereby activating G2/M checkpoint-associated stress-response genes and attenuating pathological cardiac remodeling in HFpEF.

### WEE1 inhibition ameliorates HFpEF phenotype in TGM2-deficient mice

TGM2-mediated histone serotonylation was associated with transcriptional activation of G2/M checkpoint-related genes in SAUNA-induced HFpEF hearts, whereas this transcriptional response was attenuated in TGM2-deficient mice. Then, to validate the functional relevance of this pathway, we performed rescue experiments using adavosertib, a pharmacological WEE1 inhibitor, in SAUNA-exposed TGM2-CKO mice (Figure 7A). Given that WEE1 acts as a critical negative regulator of G2/M transition through inhibitory phosphorylation of CDK1,^24^ we tested whether the WEE1 inhibition could restore CDK1-associated signaling and ameliorate the aggravated HFpEF phenotype caused by cardiomyocyte TGM2 deficiency. Treatment with adavosertib for 4 weeks significantly attenuated the increases in E/A and E/e’ ratios, as well as left atrial volume in SAUNA-exposed TGM2-CKO mice compared to vehicle-treated TGM2-CKO mice, although adavosertib did not affect systolic blood pressure, serum creatinine, or urinary albumin levels (Figure 7B-7E and Supplementary Figure 9). Furthermore, adavosertib treatment improved running distance and reduced the lung wet-to-dry ratio, kidney weight, as well as cardiac remodeling marker *Nppa* mRNA expression in SAUNA-exposed TGM2-CKO mice compared with vehicle-treated TGM2-CKO mice (Figure 7F-7H). In contrast, *Cdk1* and *Ccna2* mRNA levels were not changed between adavosertib- and vehicle-treated TGM2-CKO mice (Figure 7I), suggesting that adavosertib modulates this pathway primarily through post-transcriptional regulation of CDK1 activity rather than transcriptional induction of G2/M checkpoint–related genes. Adavosertib did not significantly alter diastolic function in SAUNA-exposed WT mice. These findings suggest that impaired G2/M checkpoint–associated signaling contributes to the HFpEF phenotype in TGM2-CKO mice and that pharmacological restoration of downstream CDK1-associated signaling partially rescues SAUNA-induced diastolic dysfunction.

### Clinical impact of circulating serotonin and cardiomyocyte H3K4me3Q5ser in patients with HFpEF

Finally, we sought to establish the clinical relevance of circulating serotonin (5-HT) and histone serotonylation in HFpEF patients. We found that circulating 5-HT levels were negatively correlated with B-type natriuretic peptide levels and tricuspid regurgitation pressure gradient (Supplementary Table 1 and 2). To evaluate the prognostic impact of circulating 5-HT, patients were stratified into two groups based on the median 5-HT levels (90.5 ng/mL). Patients with lower circulating serotonin levels had significantly lower event-free survival from the composite cardiac event of cardiac death and worsening heart failure over a median follow-up of 725 days (Figure 8A). Multivariable Cox proportional hazard analysis showed that low 5-HT levels were independently associated with a higher risk of cardiac events, with a hazard ratio of 3.25 (Figure 8B and Supplementary Table 3). Furthermore, histological analysis of myocardial biopsy specimens showed that patients with higher circulating 5-HT levels exhibited greater cardiomyocyte H3K4me3Q5ser intensity (Figure 8C and 8D). Cardiomyocyte H3K4me3Q5ser intensity was positively correlated with circulating 5-HT levels in patients with HFpEF (Figure 8E), suggesting that systemic serotonin availability may be associated with myocardial histone serotonylation status and clinical outcomes in HFpEF.

## Discussion

This study is the first to demonstrate that TGM2-mediated histone serotonylation represents a novel epigenetic cardioprotective mechanism in HFpEF. Cardiomyocyte-specific TGM2 deficiency exacerbated HFpEF-associated phenotypes, including impaired diastolic cardiac function, reduced exercise capacity, increased pulmonary congestion, and defective relaxation kinetics at the single-cardiomyocyte level. Mechanistically, under HFpEF conditions, histone serotonylation preferentially regulated genes associated with G2/M checkpoint signaling. Pharmacological inhibition of WEE1 ameliorated the HFpEF phenotypes observed in TGM2-deficient mice.

Given that HFpEF is increasingly viewed as a clinically heterogeneous syndrome rather than a single entity, recent efforts have focused on classifying patients into distinct phenogroups.^25^ The SAUNA model employed in the present study particularly mimics hypertension- and chronic kidney disease-associated HFpEF phenogroups,^26^ and our findings provide novel insights into the underlying regulatory mechanism. Although serotonin has been extensively studied for decades, its significance in the cardiovascular system remains incompletely understood. The serotonin 2B receptor has been shown to be essential for cardiac development^11^ while pharmacological antagonism of the serotonin 2B receptor suppresses post–myocardial infarction cardiac remodeling.^12^ However, these studies have focused primarily on intracellular signaling mediated by membrane-bound serotonin receptors acting as neurotransmitters, while the epigenetic roles of serotonin within the nucleus have largely been overlooked.

Histone serotonylation has been discovered as a distinct chemical modification of histones.^9^ Functionally, this modification has been shown to promote neuronal differentiation and has been implicated in neuropsychiatric disorders, including major depressive disorder as histone serotonylation enhances transcription of genes involved in neuronal activity and synaptic signaling.^9,10^ Histone serotonylation is partly mediated by intracellular serotonin uptake via the serotonin transporter, as shown by us and others, administration of high concentrations of serotonin increases H3K4me3Q5ser levels in cardiac and neuronal cells.^9^ Despite the relatively low abundance of serotonin in the heart, even modest alterations in intracellular serotonin levels and histone serotonylation may regulate cardiomyocyte gene expression, highlighting an underappreciated epigenetic control mechanism in cardiac biology.

TGM2 functions as a catalytic enzyme responsible for histone serotonylation by covalently conjugating free monoamine donors to glutamine-containing protein substrates.^9^ Although we found that cardiac H3K4me3Q5ser levels were markedly reduced at baseline in the absence of cardiomyocyte TGM2, the histone serotonylation was not completely abolished. This residual H3K4me3Q5ser may reflect contributions from non-cardiomyocyte cell types and/or compensatory mechanisms mediated by other enzymes. Bidirectional histone monoaminylation at Gln5 has been recently reported,^27^ but no enzyme responsible for removing the serotonylation has yet been identified. Histone serotonylation may represent a relatively long-lasting modification under pathophysiological conditions. Nevertheless, it remains possible that serotonylation is dynamically regulated and may be reversed by yet unidentified enzymes as part of an epigenetic regulatory mechanism. Previous studies examining cardiac phenotypes resulting from TGM2 overexpression or knockout have yielded conflicting results.^28–30^ By utilizing a cardiomyocyte-specific conditional TGM2 knockout model, our study overcomes a major limitation of prior investigations of systemic and extracardiac effects of TGM2 deficiency. Furthermore, the role of cardiomyocyte TGM2 may differ between distinct heart failure phenotypes, such as HFpEF and HFrEF. TGM2 may also exert other functions in the cytoplasm,^31^ although these roles remain to be fully elucidated.

We identified G2/M checkpoint signaling as a key target of histone serotonylation in HFpEF. Although adult cardiomyocytes are generally considered terminally differentiated, components of the G2/M checkpoint have been implicated in non-proliferative functions. Recent studies have shown that deficiency of WEE1 or pharmacological WEE1 inhibition attenuated angiotensin II-induced cardiac remodelling in mice.^32^ Thus, in cardiomyocytes, the G2/M checkpoint may function primarily as a stress-response mechanism, rather than a regulator of cell division. Further studies are needed to determine how G2/M checkpoint–associated signaling regulates cardiomyocyte relaxation kinetics under HFpEF conditions.

Modulating the TGM2–histone serotonylation pathway or its downstream G2/M checkpoint-associated signaling in a heart tissue–specific manner may represent a novel therapeutic strategy for HFpEF. In addition, assessment of cardiomyocyte histone serotonylation or circulating 5-HT levels may help identify a subset of patients with HFpEF who could benefit from personalized pathway-targeted interventions. Although pharmacological WEE1 inhibition ameliorated HFpEF phenotypes in TGM2-deficient mice, further studies are required to determine the safety, specificity, and clinical applicability of targeting this pathway in patients with HFpEF.

The limitation of the present study is that the SAUNA model was employed as a murine model of HFpEF but the generalizability of our findings to other HFpEF models or sex-specific differences remains to be determined. In the clinical study, the single-center design, relatively small sample size, and short follow-up period limit causal inference and warrant validation in larger studies with longer follow-up.

In conclusion, TGM2-mediated histone serotonylation is cardioprotective and may represent a novel therapeutic target for HFpEF.

## Methods

A comprehensive description of the materials and methods is available in the supplemental material. The data that support the findings of this study are available from the corresponding author upon reasonable request.

### Animals

B6.129S1-Tgm2tm1Rmgr/J (TGM2 flox, stock number #024694) were obtained from Jackson Laboratory. The mice carrying the *Tgm2* flox allele were crossed with transgenic mice expressing α-myosin heavy chain promoter-driven Cre recombinase^33^ to generate cardiomyocyte-specific TGM2 knockout mice. C57BL/6 mice were obtained from Jackson Laboratory Japan. Mice were housed with food and water ad libitum during 12-h light/12-h dark cycles (light, 7:00–19:00; dark, 19:00–7:00), and ambient temperature (21.5 °C) and humidity (55 ± 10%) were monitored.^34^ All animal studies were reviewed and approved by Fukushima Medical University Animal Research Committee (approval number, 2023083).

### SAUNA protocol to induce HFpEF

To induce HFpEF, mice were subjected to the SAUNA protocol.^13,14,15^ Mice were anesthetized by intraperitoneal administration of medetomidine (0.3 mg/kg), midazolam (4.0 mg/kg), and butorphanol (5.0 mg/kg). After achieving a surgical depth of anesthesia, mice underwent unilateral nephrectomy and were then exposed to a continuous infusion of d-aldosterone (0.30 µg/h; Sigma-Aldrich) via osmotic minipumps (ALZET micro-osmotic pump MODEL 1004, DURECT corporation) placed into the subcutaneous space, along with salty (1% NaCl) drinking water for 4 weeks. Control mice were subjected to sham surgery instead of unilateral nephrectomy and received continuous infusion of saline with normal drinking water.

### Echocardiographic assessment

Transthoracic echocardiography was carried out using a Vevo 2100 high-resolution imaging system (VisualSonics Inc.) equipped with a 40-MHz transducer.^35^ Parasternal short-axis M-mode images at the level of the papillary muscles were acquired in conscious mice, and measurements were averaged across at least four consecutive cardiac cycles. Fractional shortening was calculated as 100 × (LVIDd − LVIDs) / LVIDd, where LVIDd and LVIDs denote end-diastolic and end-systolic LV internal dimensions, respectively. LV mass was derived using the formula 1.05 × {[(LVIDd + IVSd + LVPWd)^3^ − (LVIDd)^3^]}, with IVSd and LVPWd representing diastolic interventricular septal and posterior wall thicknesses. For Doppler measurements, mice were lightly anesthetized with isoflurane (0.5–1.5%) in the supine position, with heart rate adjusted to approximately 400 beats per minute. Transmitral inflow was recorded by pulsed-wave Doppler from an apical four-chamber view, with the sample volume positioned at the tips of the mitral valve leaflets and aligned parallel to flow. Peak early (E) and late (A) diastolic velocities were measured, and the E/A ratio was calculated. Recordings with fused E and A waves were excluded from analysis. Tissue Doppler imaging was used to measure early diastolic mitral annular velocity (e′) at the mitral annulus, and E/e’ ratio was calculated. Left atrial volume was measured by tracing the left atrial endocardial border at end-systole.

### Blood pressure measurements

The systolic, mean, and diastolic blood pressures, as well as heart rate, were measured by the tail-cuff method using a programmable sphygmomanometer (BP-98A-L, Softron, Tokyo, Japan) in the conscious mice.^36^ The data were averaged over 5 consecutive measurements.

### Measurement of serum creatinine, blood urea nitrogen, and urinary albumin

Serum creatinine, blood urea nitrogen, and urinary albumin levels were measured by Oriental Yeast Co., ltd.

### Treadmill exercise testing

Exercise capacity in mice was evaluated using a motor-driven treadmill system (MK-685, Muromachi Kikai Co) after a three-day period of treadmill habituation.^37^ The treadmill was set at a 20° incline. Each mouse began running at an initial speed of 5 m/min for warm-up, after which the speed was progressively increased to 14 m/min. This maximum speed was maintained until the animal reached exhaustion. Exhaustion was defined as the inability to resume running within 10 s after direct contact with the electrical stimulus grid. Total running duration was measured, and the running distance was calculated accordingly.

### Western blot analysis

Snap frozen mouse heart samples or cultured cells were initially homogenized in lysis buffer (Cell Lysis Buffer, Cell Signaling Technology) containing protease inhibitor cocktail (BD Biosciences). Protein concentrations were quantified using the BCA Protein Assay Kit (Pierce, Thermo Fisher Scientific). Equal amounts of protein were resolved by SDS-polyacrylamide gel electrophoresis and transferred to nitrocellulose membranes (Amersham Protran Premium 0.2 NC, GE Healthcare). After the membrane was blocked using 5% skim milk for 1 h, membranes were incubated overnight at 4°C with the following specific antibodies; H3K4me3Q5ser (1:1000, ABE2580, Merck KGaA), TGM2 (1:1000, 3557, Cell signaling technology), GAPDH (1:1000, 60004-1-Ig, Proteintech), Histone H3 (1:1000, 9715, Cell signaling technology), followed by the appropriate IRDye 680 or IRDye 800 secondary antibodies (1:20000, 925-68070, 925- 68071, 925-32210, 925-32210, LI-COR Biosciences). Fluorescent immunoreactive bands were detected by an Odyssey CLX imaging system (LI-COR Biosciences). Optical densities of individual bands were analyzed using ImageJ software (National Institutes of Health) or Image Studio software (LI-COR Biosciences).

### Isolation of nuclear fraction

Nuclear fraction was isolated from heart tissues using differential centrifugation.^38^ Samples were homogenized in 250 mM sucrose, 50 mM Tris-HCl, pH 7.4, 5 mM MgCl2 and centrifuged at 800 × g for 15 min at 4°C. The nuclear pellet was washed and resuspended in nuclear extraction buffer that contains 20 mM HEPES, pH 7.9, 1.5 mM MgCl_2_, 0.5 M NaCl, 0.2 mM EDTA, 20% glycerol for subsequent analyses.

### Pseudobulk normalization and donor-level aggregation of snRNA-sequencing data

Donor-level comparisons in the human single-nucleus RNA-sequencing dataset SCP1303^16^ were performed using a pseudobulk approach. For each donor, raw TGM2 unique molecular identifier (UMI) counts were summed across cardiomyocyte nuclei, normalized to the total UMI counts of the corresponding cardiomyocyte nuclei, and multiplied by 10□. The resulting values were expressed as counts per million UMIs and used for donor-level comparisons.

### Histological analysis and fluorescent immunohistochemistry of murine hearts

Murine LV samples were fixed in 4% paraformaldehyde solution for paraffin embedding or embedded in the O.C.T. compound (Tissue-Tek). Paraffin-embedded sections (2 μm thickness) were stained with hematoxylin-eosin or Elastica-Masson for assessment of myocardial fibrosis.^35^ The myocardial fibrosis fraction was measured using ImageJ software (National Institutes of Health). For the assessment of cardiomyocyte cross-sectional area, paraffin-embedded tissue sections were incubated with wheat germ agglutinin conjugated to Alexa Fluor 488 (Thermo Fisher Scientific) and mounted with DAPI-containing medium (Fluoro-Gel II with DAPI, Electron Microscopy Sciences).^35^ Cardiomyocyte cross-sectional area was quantified from more than 100 cells using ImageJ software. Fluorescent immunohistochemistry was carried out on O.C.T. compound-embedded LV sections (6 μm thickness). All microscopic images were obtained with a Keyence BZ-X700 microscope and processed with the BZ II Viewer software (Keyence).

### Reverse transcription-quantitative polymerase chain reaction (RT-qPCR)

Total RNA was extracted from cultured cells or mouse hearts using TRIzol reagent (Thermo Fisher Scientific) according to the manufacturer’s protocol. cDNA was synthesized using ReverTra Ace qPCR RT Master Mix (Toyobo Co., Ltd.). Quantitative PCR was performed using THUNDERBIRD SYBR qPCR Mix (Toyobo Co., Ltd.) in a CFX Connect real-time PCR System (Bio-Rad) with Bio-Rad CFX Manager 3.1 software (Bio-Rad). The mRNA expression levels of *Nppa*, *Col3a1*, *Cdk1*,and *Ccna2* were analyzed using the 2^-ΔΔCt^ method. Data were normalized to *18S rRNA* in mouse or *Gapdh* in rat and expressed as a fold increase of the control group. Primer sequences are described in the Supplementary Table 4.

### Transmission electron microscopy

After mice were anesthetized by intraperitoneal administration of medetomidine (0.3 mg/kg), midazolam (4.0 mg/kg), and butorphanol (5.0 mg/kg), mice were perfused with heparinized saline (1 U/mL). The hearts were then perfusion-fixed with ice-cold 0.1 M phosphate buffer (pH 7.4) containing 2% glutaraldehyde and 2% paraformaldehyde. Samples were excised and post-fixed using the reduced-osmium method, embedded in Epon812, and cut into ultrathin sections. After staining with uranyl acetate and lead citrate, the sections were examined with a transmission electron microscope (JEOL, JEM1400EX).

### Imaging mass spectrometry for 5-HT detection

Mass spectrometry imaging was performed to visualize 5-HT distribution in mouse heart sections using TMPy derivatization-based imaging MS.^40^ Cryo-embedded heart sections were mounted onto indium tin oxide–coated glass slides, derivatized with 2,4,6-trimethylpyrylium tetrafluoroborate, and sprayed with CHCA matrix solution. Imaging mass spectrometry was performed in positive ion mode using a MALDI mass spectrometer (Bruker Daltonics). Spectra were acquired at a spot-to-spot distance of 100 μm with 200 laser shots per spot. TMPy-labeled 5-HT was detected at m/z 280.205, and ion images were reconstructed using FlexImaging 4.0 software (Bruker Daltonics) with a mass window of m/z ± 0.050. Peak intensities were normalized to the total ion current.

### Cell shortening and relaxation kinetics in isolated cardiomyocytes

Adult mouse cardiomyocytes were isolated from mice using a Langendorff-free method.^41^ After the mice were anesthetized with 5% isoflurane, the ascending aorta was clamped, and the heart was quickly excised. The left ventricle was perfused with EDTA buffer from the apex, followed by digestion buffer containing collagenase type 2 (1 mg/mL, Worthington Biochemical Corporation) and protease type XIV (50 μg/mL, Sigma-Aldrich) at 37°C. After stepwise calcium reintroduction, rod-shaped cardiomyocytes were plated onto laminin (20 μg/mL, Sigma-Aldrich)-coated plates.

Isolated cardiomyocytes were seeded on 35-mm μ-ibiTreat high imaging dishes (Ibidi). Cells were electrically field-stimulated at 20 V and 1 Hz, and changes in cell length were recorded using a CMOS camera (CHU30, Shodensha Co., Ltd.). Cell shortening and relaxation kinetics were analyzed from time-series images at the single-cell level.^17,18^ Cell shortening was calculated as 100 × (resting cell length − minimal cell length during contraction) / resting cell length.

### Cell culture and isolation of neonatal rat cardiomyocytes (NRCM)

H9c2 rat embryonic cardiac myoblasts were purchased from ATCC (CRL-1446). The cells were cultured in Dulbecco’s modified Eagle’s medium (DMEM, FUJIFILM Wako) supplemented with 10% fetal bovine serum, 100 μg/mL of streptomycin, and 100 IU/mL of penicillin at 37 °C in the presence of 5% CO_2_. NRCM was isolated from hearts of 1- to 2-day-old Sprague-Dawley rats obtained from CLEA Japan with enzymatic dissociation by collagenase type II (Worthington) and were cultured in DMEM containing 20% fetal bovine serum, 10% penicillin-streptomycin-glutamine, and 100 μM BrdU (Sigma) for 24 h. The media were changed into serum-free DMEM containing 10% penicillin-streptomycin-glutamine, 100 μM BrdU (Sigma), 1 μg/mL insulin (Sigma), and 5 μg/mL transferrin (Sigma).

### Administration of finerenone

Finerenone (BAY 94-8862) was purchased from Selleck Chemicals and administered orally once daily at a dose of 10 mg/kg body weight for 4 weeks in SAUNA-exposed WT mice.

### CUT&RUN library preparation and sequencing

CUT&RUN was performed using CUT&RUN assay kit (86652, Cell signaling technology) following the manufacturer’s protocol.^19^ Hearts were initially homogenized and 5 mg of tissue bound to concanavalin A-coated magnetic beads. Chromatin was incubated with 2 μL of anti-H3K4me3Q5ser antibody (ABE2580), and 50 pg of spike-in control DNA provided with the kit was added to each sample for normalization. DNA was purified by DNA Purification Buffers and Spin Columns (14209, CST). Libraries were prepared from CUT&RUN DNA using the NEBNext Ultra II DNA Library Prep Kit for Illumina at Rhelixa Inc. Sequencing was performed on an Illumina NovaSeq 6000 platform as 150-bp paired-end reads (PE150). Approximately 6 Gb of sequencing data were generated per sample, corresponding to approximately 40.0 million reads per sample, or 20 million read pairs per sample. The sequencing data obtained were used for downstream peak calling and enrichment analyses.

### CUT&RUN data analysis

Normalized signal tracks were generated from mapped BAM files using bamCoverage in deepTools v3.5.6. Read coverage was normalized using the RPGC method with an effective mouse genome size of 2,652,783,500 bp for mm10. Reads were extended to 150 bp and centered, and coverage was calculated using a bin size of 10 bp. PCR duplicates, reads with mapping quality scores below 30, and genomic regions included in the ENCODE mm10 blacklist v2 were excluded from the analysis. The resulting bigWig files were used for genome browser visualization, metagene profiling, and heatmap generation.

Metagene profiles and heatmaps were generated using computeMatrix, plotProfile, and plotHeatmap in deepTools. Genomic coordinates were obtained from the UCSC mm10 known gene annotation using the TxDb.Mmusculus.UCSC.mm10.knownGene R package. For pathway-focused visualization, G2/M checkpoint-related genes were defined using the mouse Hallmark gene set collection retrieved with the msigdbr R package. For GSEA, differential enrichment of H3K4me3Q5ser was analyzed using DiffBind v3.18.0 with aligned BAM files and stringent consensus peak sets as input. Consensus peaks were defined as peaks detected in two biological replicates within each group. Peak boundaries defined by SEACR were retained without re-centering, and peaks with a total read count below 10 across all samples were removed. Raw read counts were normalized using the DESeq2 method implemented in DiffBind, and differential binding was assessed for the indicated comparisons. Differentially enriched peaks were annotated to the nearest transcription start sites within ±3 kb using ChIPseeker and the TxDb.Mmusculus.UCSC.mm10.knownGene database. When multiple promoter-associated peaks were assigned to the same gene, the peak with the lowest P value was selected as the representative peak. Pre-ranked gene set enrichment analysis was then performed using clusterProfiler. Genes were ranked by a P-value-weighted metric calculated as sign(fold change) × −log10(P value). The MSigDB Hallmark gene set collection was used as the reference, and gene sets containing fewer than 50 genes were excluded. Pathways with an FDR q value < 0.25 were considered potentially enriched and prioritized for downstream interpretation.

### RNA sequencing

Total RNA was isolated from 20 mg of mouse heart tissue using TRIzol reagent (Thermo Fisher Scientific) and purified with a DNA-free DNA Removal Kit (Thermo Fisher Scientific).^35^ RNA concentrations were measured using a NanoDrop 2000/2000c Spectrophotometer (Thermo Fisher Scientific). One microgram of purified total RNA per sample was submitted to BGI Japan for library preparation and sequencing. RNA-seq libraries were prepared with a poly(A)-selected, strand-specific library preparation method and sequenced using BGISEQ-500 (Beijing Genomics Institution) to generate 150-bp paired-end reads. Low-quality reads and adapter-contaminated reads were filtered using SOAPnuke. Clean reads were aligned to the mouse reference genome, Mus_musculus_10090.UCSC.mm10.v2201, using HISAT2, and gene expression was quantified using RSEM. Raw count data obtained from the BGI Dr. Tom systems were used for downstream analyses. Count data were normalized using the variance-stabilizing transformation in DESeq2. For heatmaps, normalized expression values were row-scaled to generate Z-scores. Samples were ordered by experimental group, and genes were hierarchically clustered using Ward’s method. Interaction effects estimated from the DESeq2 model were visualized as scatter plots.

### Gene set enrichment analyses (GSEA)

GSEA was conducted using the clusterProfiler and fgsea packages. Genes were ranked by their log2 fold change values calculated from the DESeq2 analysis. The MSigDB Hallmark gene set collection (h.all.v2023.2.Mm) was used as the reference database. GSEA was performed for four specific comparison patterns to isolate the effects of the disease and the rescue effect of TGM2-KO. Significance was defined by a false discovery rate (FDR) < 0.25 and a nominal p-value < 0.05. For the prioritized pathways, enrichment plots (Running ES plots) were generated. The results of the pathway analysis were visualized using bubble charts to compare the normalized enrichment scores (NES) across different experimental conditions.

### Leading edge analysis and protein-protein interaction network

The protein-protein interaction network was constructed for these leading-edge genes using the STRING database (v11.5) for Mus musculus.^42^ An interaction score threshold of 900 (Highest confidence) was applied to ensure high stringency, and the resulting network was analyzed using the igraph, tidygraph, and ggraph packages. The network layout was optimized using the Kamada-Kawai (KK) force-directed algorithm. To identify key hub genes within the network, topological metrics including Degree centrality and Betweenness centrality were calculated. Hub genes were defined based on high Degree scores and visualized with node sizes and colors proportional to their topological importance. All visualizations were produced using ggplot2 and ggpubr.

### Construction of DNA plasmid and site-directed mutagenesis

To generate a GFP-TGM2 plasmid, mouse *Tgm2* (NM_009373.3) cDNA was obtained by RT-PCR from RNA from C57BL/6J mouse heart. The cDNA was amplified by PCR using the following primers: forward, 5’-AAAGAATTCCATGGCAGAGGAGCTGCTCCTGGAGA-3’; reverse, 5’-AAAGGTACCTTAGGCCGGGCCGATGATAACATTCC-3’. The DNA fragment was subcloned into pAcGFP1-C1 vector (632470, Clontech) at EcoRI and KpnI sites. To generate mutant GFP-TGM2 plasmids, PCR-based site-directed mutagenesis was performed using PrimeSTAR MaxDNA polymerase (Takara Bio). Mutant GFP-TGM2 constructs containing N398A/D400A substitutions and lacking amino acids 596–601 (PKQNRK) was generated. The primers for mutagenesis were as follows; for the N398A/D400A mutant, forward, 5’-GTCGCCGCTGCAGTGGTGGACTGGATCCGG-3’; reverse, 5’- CACTGCAGCGGCGACCTCGGCAAACACGAA-3’; forward, 5’-TGGGAGAACTGGTGGCTGAAGGTGTC-3’; and for the Δ596–601 (PKQNRK) mutant, reverse, 5’- CCACCAGTTCTCCCAGGACCCGGAT, -3’. All plasmid constructs were verified by restriction digestion and DNA sequencing.

### Transfection with plasmid DNA

H9c2 cardiac myocytes were transfected using Lipofectamine 3000 (Thermo Fisher Scientific) according to the manufacturer’s instructions.

### Immunofluorescence and time-lapse live-cell imaging

H9c2 myocytes were placed on 35-mm μ-ibiTreat high imaging dishes (Ibidi). Cells were fixed with 4% paraformaldehyde, and then mounted with DAPI-containing mounting media.

### WEE1 inhibitor administration

Adavosertib (MK-1775, Selleck Chemicals) was administered orally to mice once daily at a dose of 50 mg/kg body weight for 4 weeks in SAUNA-exposed mice.

### Human HFpEF cohort

The human HFpEF cohort study enrolled consecutive 240 hospital-admitted patients presenting with symptomatic stage C/D HFpEF at Fukushima Medical University Hospital between November 2017 and March 2022. We excluded patients with cardiac disease requiring surgical intervention, end-stage renal disease with hemodialysis, and chronic lung disease requiring oxygen therapy. Serum 5-HT levels were measured by Serotonin ELISA kit BA-E-8900R (ImmuSmol). Clinical information, including demographic data, laboratory data, echocardiographic analysis was collected as part of standard clinical practice.^43,44^ All patients were followed up by reviewing medical records at the same hospital or referral hospitals for composite cardiac events that include cardiac death, unplanned re-hospitalization due to worsening heart failure.^44^ Cardiac death was defined as death from heart failure, serious ventricular arrhythmia that includes ventricular fibrillation and ventricular tachycardia, or sudden cardiac death. Worsening heart failure was defined as symptoms and signs of heart failure that required urgent therapy and resulted in hospitalization. All endpoints were reviewed by more than two independent cardiologists on the committee. For the immunohistochemical study, 18 patients with HFpEF who underwent endomyocardial biopsy were enrolled between December 2014 and February 2022. The study protocol was approved by the Ethics Committee of Fukushima Medical University (approval numbers, 823, REC2024-161, and 2020-309). All patients provided written informed consent prior to study enrolment.

### Immunohistochemical study of human biopsy specimens

Fluorescent immunohistochemistry was performed on paraffin-embedded biopsy specimens. After deparaffinization and rehydration, the 2-μm-thick sections were stained with the following primary antibodies: H3K4me3Q5ser (ABE2580) and troponin I (ab188877, Abcam) followed by the appropriate secondary antibodies, Alexa Fluor 488–conjugated antibody (1:1000, ab150105, Abcam) and Alexa Fluor 647–conjugated antibody (1:1000, ab150159, Abcam), and mounted with DAPI-containing mounting medium (Fluoro-Gel II with DAPI, Electron Microscopy Sciences). For quantitative analysis, images were acquired systematically across the entire area of each human biopsy specimen. Fluorescence intensity was measured using ImageJ (National Institutes of Health) in at least 40 cardiomyocyte nuclei per sample. The median nuclear fluorescence intensity for each sample was used as the representative fluorescence intensity.^10^

### Statistical analysis

Continuous data are expressed as mean ± standard deviation or standard error, and skewed data are presented as median and interquartile range. Categorical variables are expressed as numbers and percentages. The Shapiro-Wilk test was used to assess normality. Comparisons between the two groups were performed using Student’s t-test or Mann-Whitney U-test. One-way ANOVA followed by multiple comparisons with the Tukey test was performed when more than two groups were evaluated. Correlations were assessed using Spearman’s correlation analysis. Categorical variables were compared using Chi-square test or Fisher’s exact test. Composite events were calculated using Kaplan-Meier analysis with the log-rank test. Univariable and multivariable Cox proportional hazards models were used to assess the association between circulating serotonin levels and cardiac events. Data were analyzed using Statistical Package for Social Sciences version 26.0 (IBM), GraphPadPrism version 8.1.2 (GraphPad Software), and R software. A value of P < 0.05 was considered statistically significant. For genome-wide analyses, P values were adjusted for multiple testing using the Benjamini–Hochberg method, and FDR q values were reported where appropriate. Gene sets with FDR q < 0.25 were considered enriched for GSEA.

## Acknowledgements

The authors thank Tomiko Miura, Tomomi Ogata, Kumiko Watanabe, Yumi Yoshihisa, and Shiori Togashi for their technical assistance; Yumiko Kurosu and Mutsuko Honda for assistance with electron microscopy procedures; Ryoma Ito and Kotaro Kodama for experimental support. This work was supported by Japan Society for the Promotion of Science KAKENHI grants 22K08107 to T.M.. Figures were generated with BioRender.com.

## Author contributions

R.O. and T.M. conceived and designed the study, performed experiments, analyzed and interpreted the data, and wrote the manuscript. Y.S., S.O., S.I., S.M., and T.Y. contributed to experimental work and data analysis. S.T. performed imaging mass spectrometry. S.W. performed transmission electron microscopy. M.O., A.Y., and T.I. provided supervision and critical scientific input. Y.T. supervised the study and approved the final version of the manuscript. All authors reviewed and approved the manuscript.

## Competing interests

TM belongs to a department supported by Fukushima Daiichi Hospital, and TY belongs to a department supported by Minamisoma City. The remaining authors have no disclosures to report.

## Data availability statement

The CUT&RUN and RNA-sequencing data generated in this study have been deposited in the DNA Data Bank of Japan Sequence Read Archive under BioProject accession code PRJDB40369. Any remaining raw data will be available from the corresponding author upon reasonable request.

