## Supplementary Material for "TGM2-mediated histone serotonylation is an epigenetic cardioprotective mechanism in HFpEF"

[1]Department of Cardiovascular Medicine, Fukushima Medical University, Fukushima, Japan. [2]Department of Community Heart Failure Medicine, Fukushima Medical University, Fukushima, Japan. [3]Department of Community Cardiovascular Medicine, Fukushima Medical University, Fukushima, Japan. [4]Faculty of Food and Agricultural Sciences, Fukushima University, Fukushima, Japan. [5]Department of Anatomy and Histology, Fukushima Medical University, Fukushima, Japan. [6]Department of Clinical Laboratory Sciences, Fukushima Medical University, Fukushima, Japan.

### Supplementary Information

#### Supplementary Figures 1-9

#### Supplementary Tables 1-4

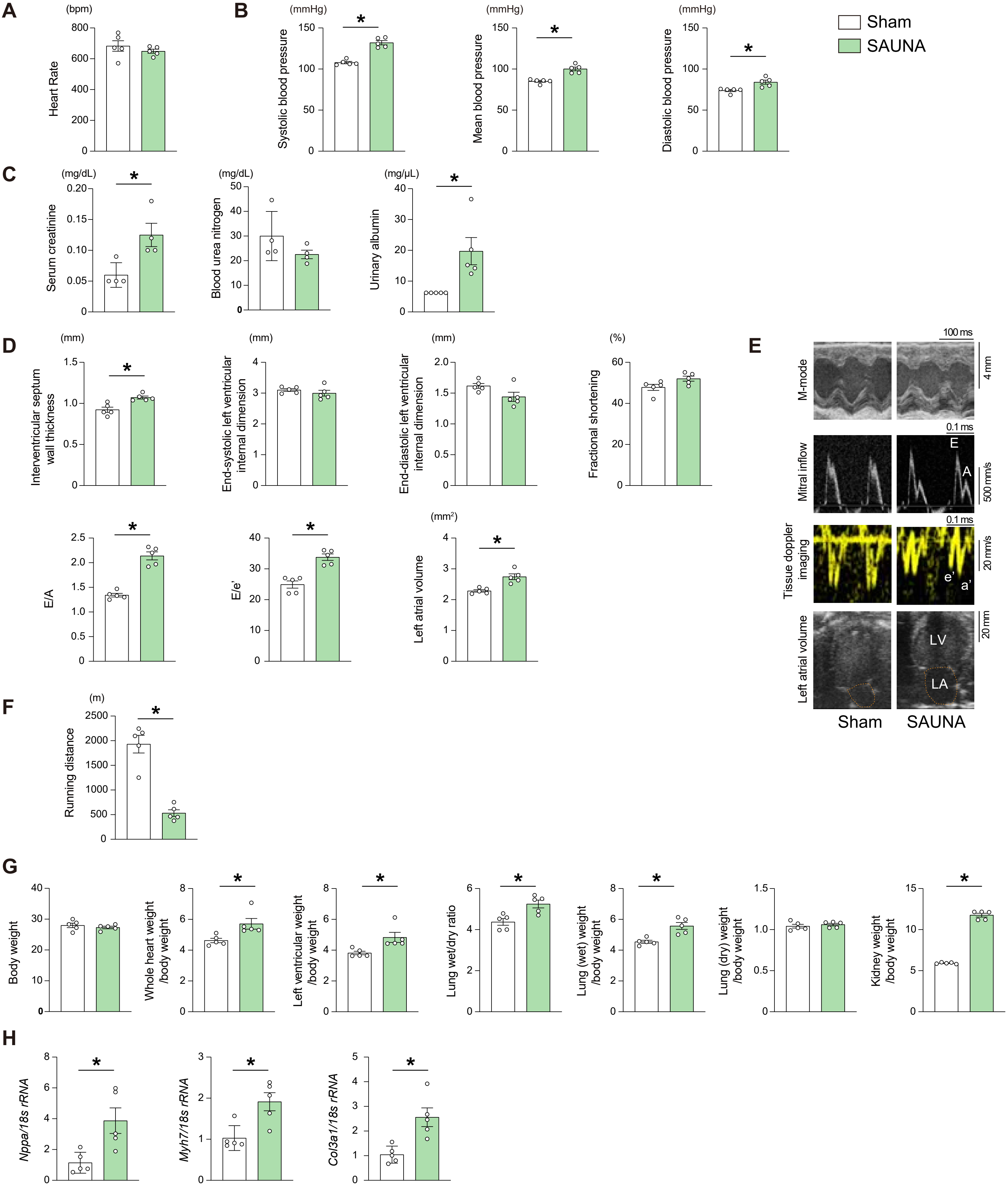


**Supplementary Figure 1. The SAUNA model recapitulates features of HFpEF.**
C57BL/6 wild-type mice were subjected to the SAUNA protocol [1% saline drinking water, unilateral nephrectomy, and continuous aldosterone infusion (0.30 μg/h)] or sham operation. **A**, Heart rate. **B**, Blood pressure. **C**, Serum creatinine and blood urea nitrogen, and urinary albumin levels. **D**, Echocardiographic parameters. **E**, Representative echocardiographic images of M-mode, pulsed-wave Doppler and tissue Doppler imaging. Peak early (E) and late (A) transmitral inflow velocities and early (e') and late (a) diastolic mitral annular velocity are shown. **F**, Running distance determined by treadmill exercise testing. **G**, Physiological parameters. **H**, Relative mRNA expression levels of *Nppa*, *Myh7*, and *Col3a1* in the heart. The data were normalized to 1*8s rRNA* levels. All data are presented as mean ± SEM. *P < 0.05 versus sham by the unpaired t test (two-sided). n=5 per group.

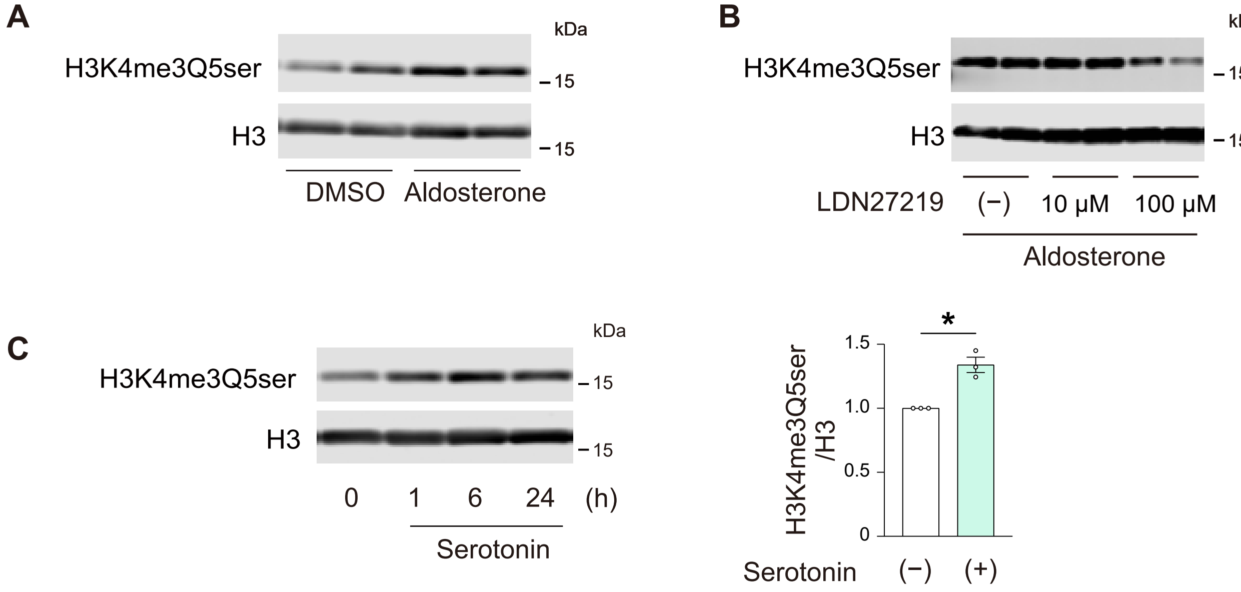


**Supplementary Figure 2. Alteration of cardiac histone serotonylation levels in cardiac myocytes.**
**A**, Representative immunoblots of H3K4me3Q5ser levels in H9c2 cardiac myocytes stimulated with aldosterone (100 nM) for 3 h. **B**, Effects of the TGM2 inhibitor LDN27219 on H3K4me3Q5ser in aldosterone-treated H9c2 cells. Cells were pretreated with LDN27219 at the indicated concentrations for 2 h, followed by stimulation with aldosterone (100 nM) for 3 h. Representative immunoblots of H3K4me3Q5ser are shown. **C**, Effects of serotonin (100 μM) on H3K4me3Q5ser in H9c2 cells for the indicated time points. Quantification of H3K4me3Q5ser levels after 24 h of serotonin stimulation (n=3). Data are presented as mean ± SEM. *P < 0.05 by two-sided unpaired t test.

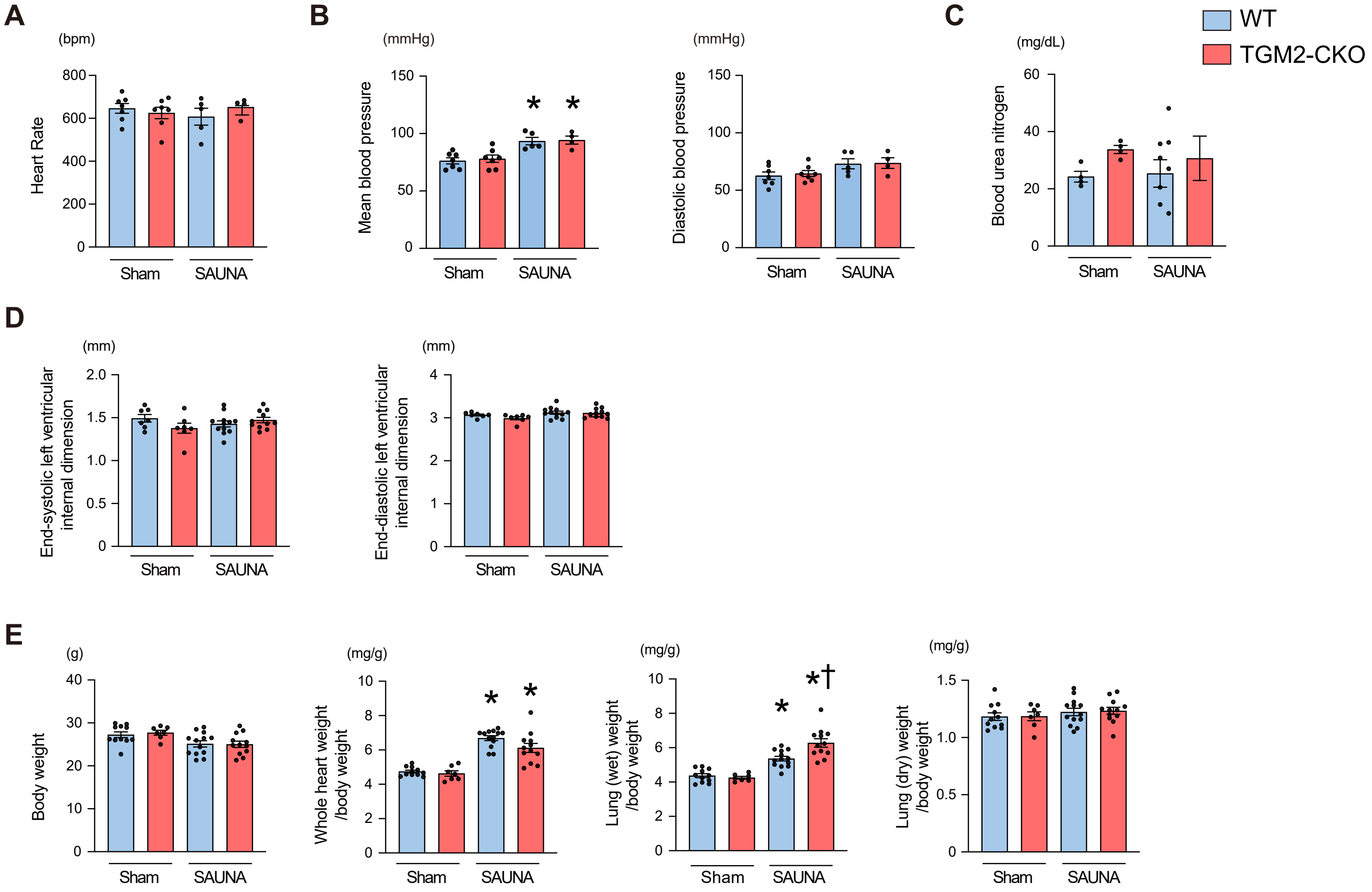


**Supplementary Figure 3. Characterization of cardiomyocyte-specific TGM2 knockout (TGM2-CKO) mice 4 weeks after SAUNA.**
**A**, Heart rate. **B**, Mean and diastolic blood pressure. **C**, Blood urea nitrogen. **D**, Echocardiographic parameters. **E**, Physiological parameters. All data are presented as mean ± SEM. *P < 0.05 versus the corresponding sham-exposed group and †P < 0.05 versus the corresponding WT mice by the one-way ANOVA with Tukey post-hoc analysis.

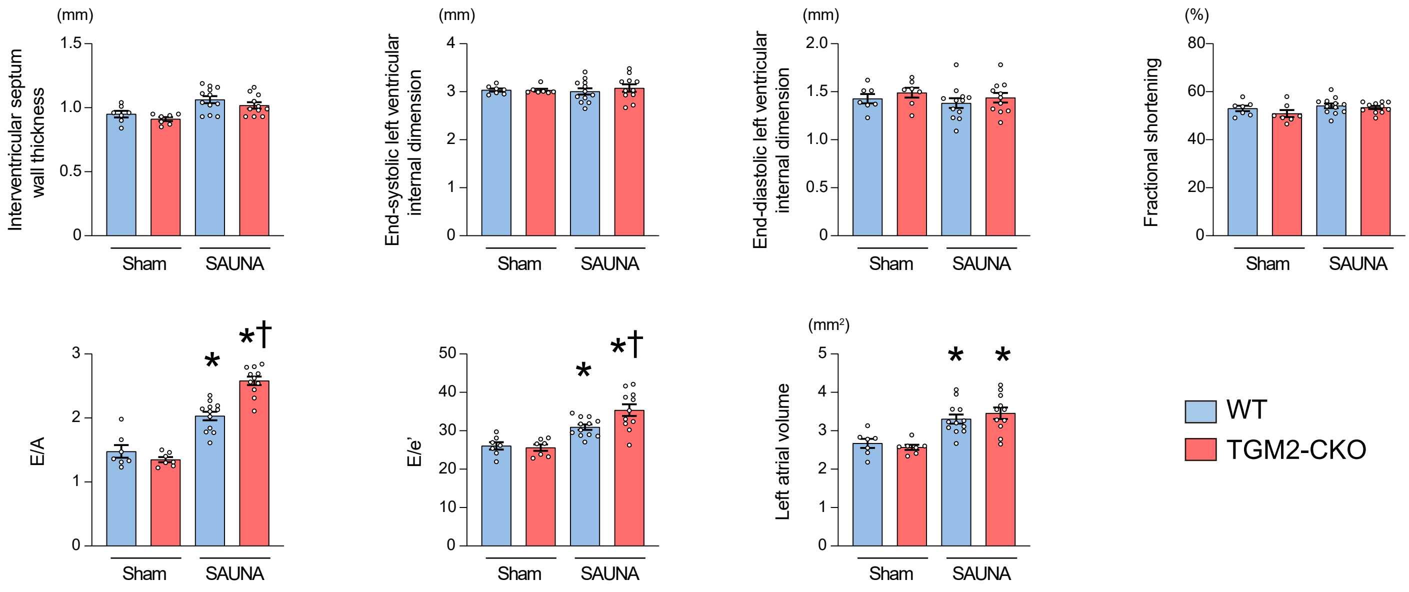


**Supplementary Figure 4. Echocardiographic parameters of cardiomyocyte-specific TGM2 knockout mice 2 weeks after SAUNA.**
All data are presented as mean ± SEM (n=7-12). *P < 0.05 versus the corresponding sham-exposed group and †P < 0.05 versus the corresponding WT mice by the one-way ANOVA with Tukey post-hoc analysis.

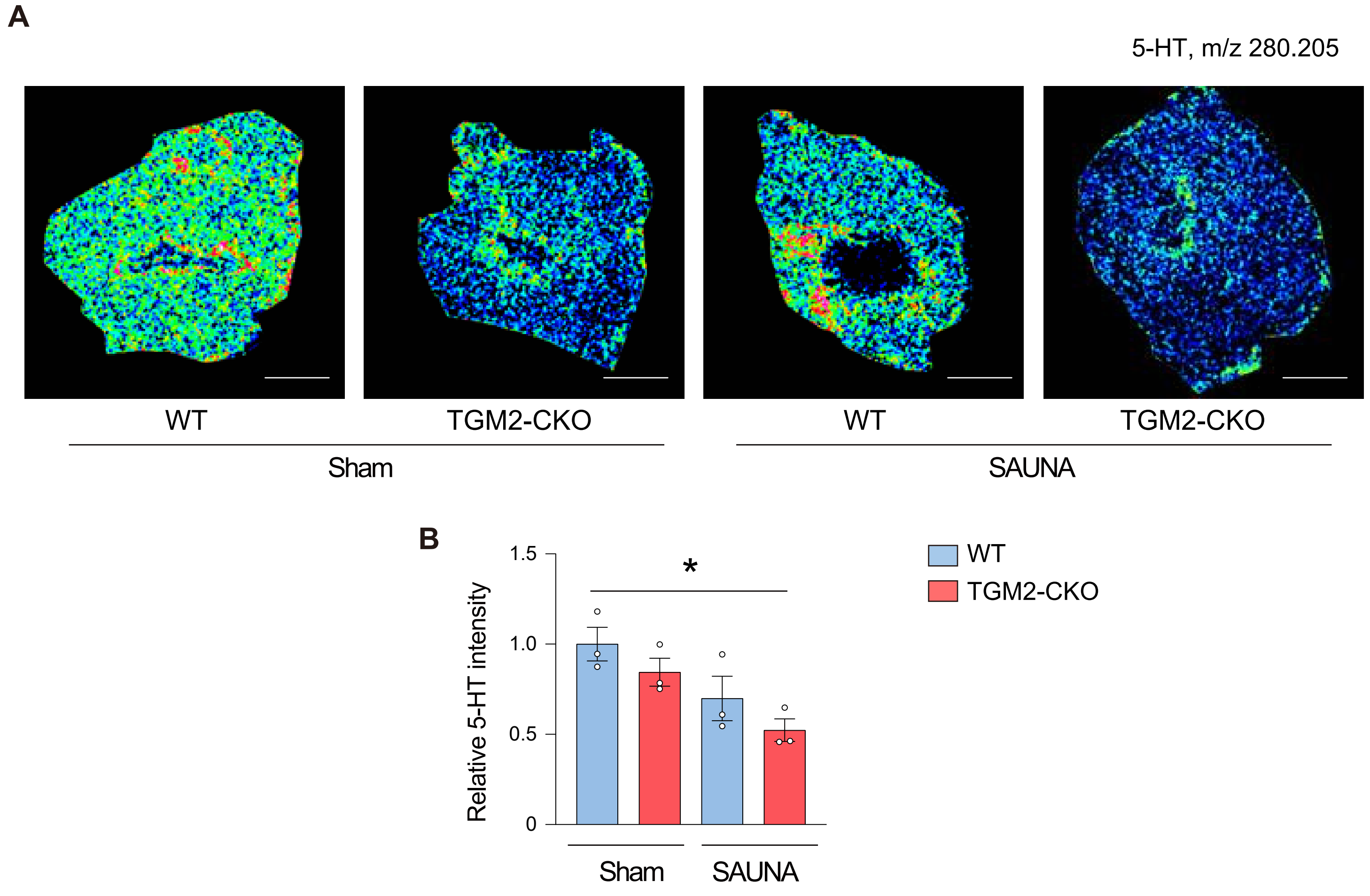


**Supplementary Figure 5. Mass spectrometry imaging of 5-HT in the left ventricle after SAUNA.**
**A**, Representative mass spectrometry images of left ventricular sections from Sham and SAUNA-exposed mice 4 weeks after SAUNA exposure. Signals were adjusted for 5-HT at m/z 280.205. Scale bars, 1 mm. **B**, Quantification of 5-HT signal intensity in the left ventricle (n=3). The peak intensity of the spectra was normalized to the total ion count and expressed as a relative ratio to Sham-exposed WT mice. Data are presented as mean ± SEM. *P < 0.05 by one-way ANOVA with Tukey post-hoc analysis.

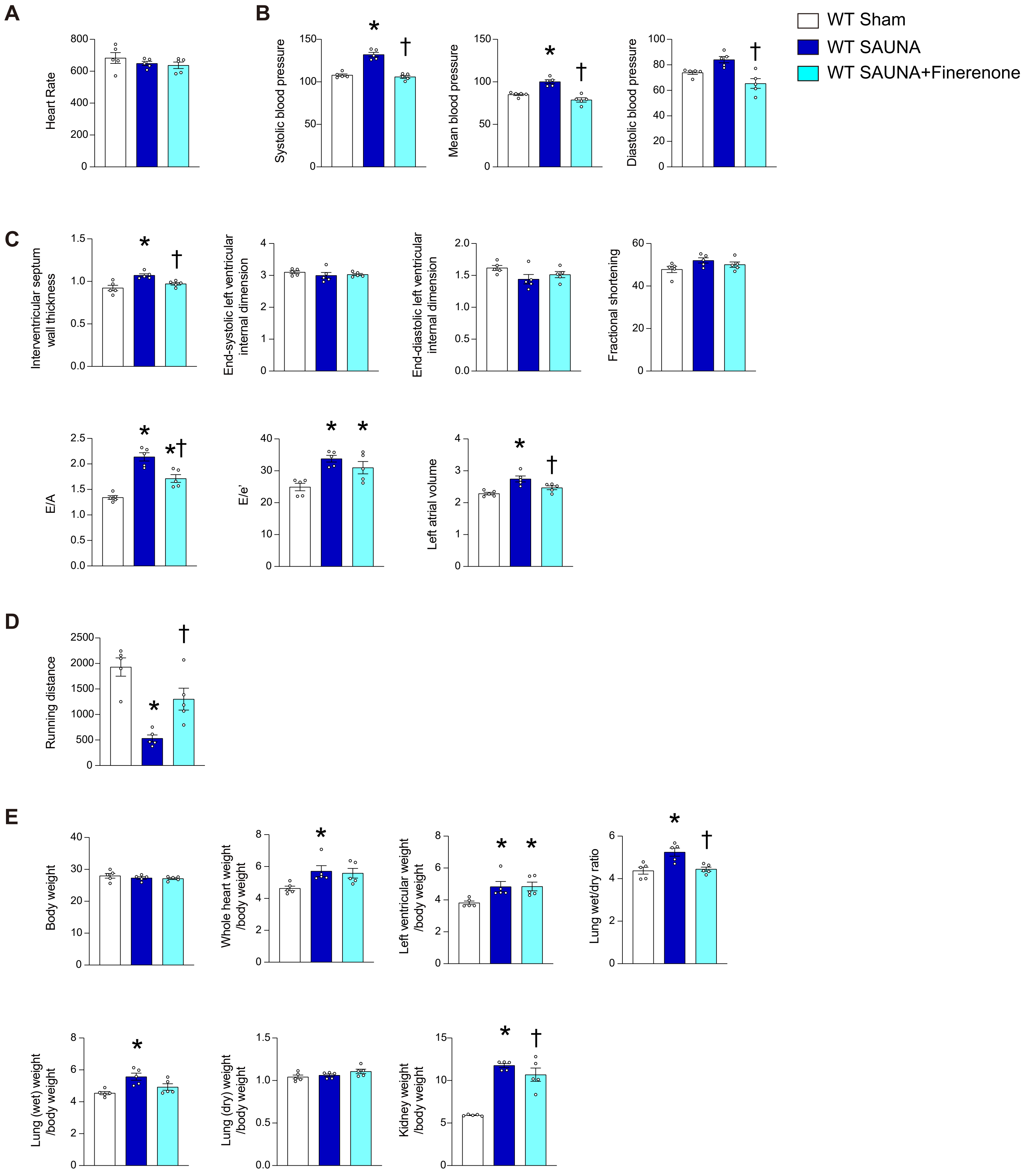


**Supplementary Figure 6. Effects of finerenone treatment in SAUNA-exposed WT mice.**
Finerenone was administered orally once daily at a dose of 10 mg/kg body weight for 4 weeks in SAUNA-exposed WT mice. **A**, Heart rate. **B**, Blood pressure. **C**, Echocardiographic parameters. **D**, Running distance determined by treadmill exercise testing. **E**, Physiological parameters. Data for WT-sham and WT-SAUNA groups are reproduced from main Figures 2 and 3. All data are presented as mean ± SEM. $^{*}P < 0.05$ versus the sham-exposed group and $^{\dagger}P < 0.05$ versus the SAUNA-exposed WT mice by the one-way ANOVA with Tukey post-hoc analysis. n=5 per group.

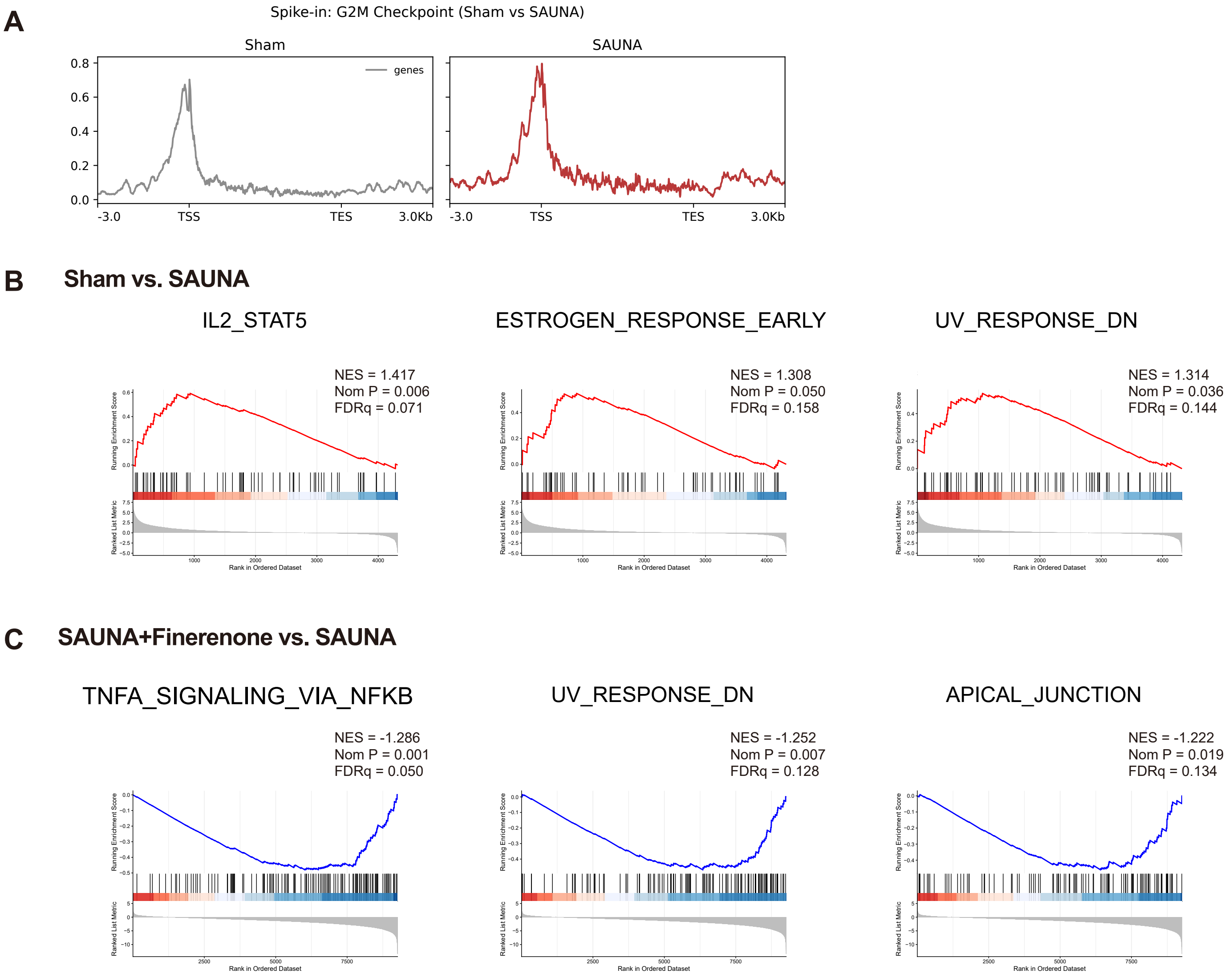


**Supplementary Figure 7. Genome-wide effects of H3K4me3Q5ser in the hearts of HFpEF.**

**A**, Metagene plots showing spike-in–normalized H3K4me3Q5ser CUT&RUN signals across G2/M checkpoint–associated genes in sham- and SAUNA-exposed hearts. Signals are displayed from 3.0 kb upstream of the transcription start site (TSS) through gene bodies to 3.0 kb downstream of the transcription end site (TES). The y-axis indicates normalized signal intensity. **B**, Gene set enrichment analysis (GSEA) plots showing pathways enriched in SAUNA-exposed hearts compared with sham hearts. **C**, GSEA plots showing gene sets negatively enriched in finerenone–treated SAUNA hearts compared with SAUNA-exposed hearts. Normalized enrichment scores (NES), nominal P values, and false discovery rates (FDR) are shown in each plot.

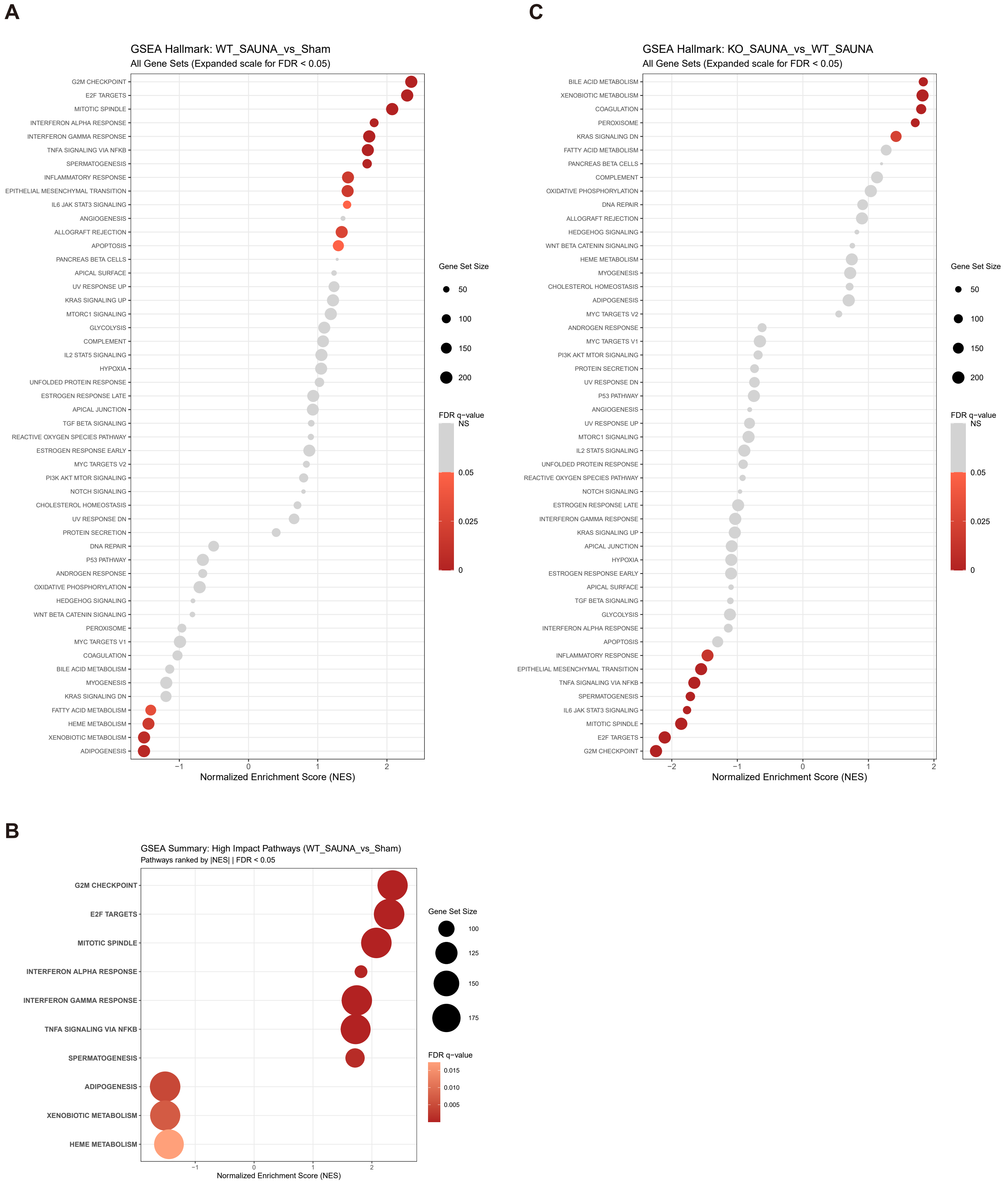


**Supplementary Figure 8. Hallmark pathway enrichment analysis of cardiac RNA-seq data.**
**A**, Bubble plot showing Hallmark pathway enrichment in WT SAUNA versus sham hearts. SAUNA exposure induced enrichment of G2/M checkpoint, E2F targets, mitotic spindle, inflammatory signaling, and interferon response pathways in WT hearts. **B**, Summary plot of significantly enriched high-impact pathways in the WT SAUNA versus sham comparison, ranked by absolute normalized enrichment score. **C**, Bubble plot showing Hallmark pathway enrichment in TGM2-CKO SAUNA versus WT SAUNA hearts. Bubble size represents gene set size, and color indicates the FDR q-value. Positive and negative normalized enrichment scores indicate enrichment in the former and latter groups of each comparison, respectively.

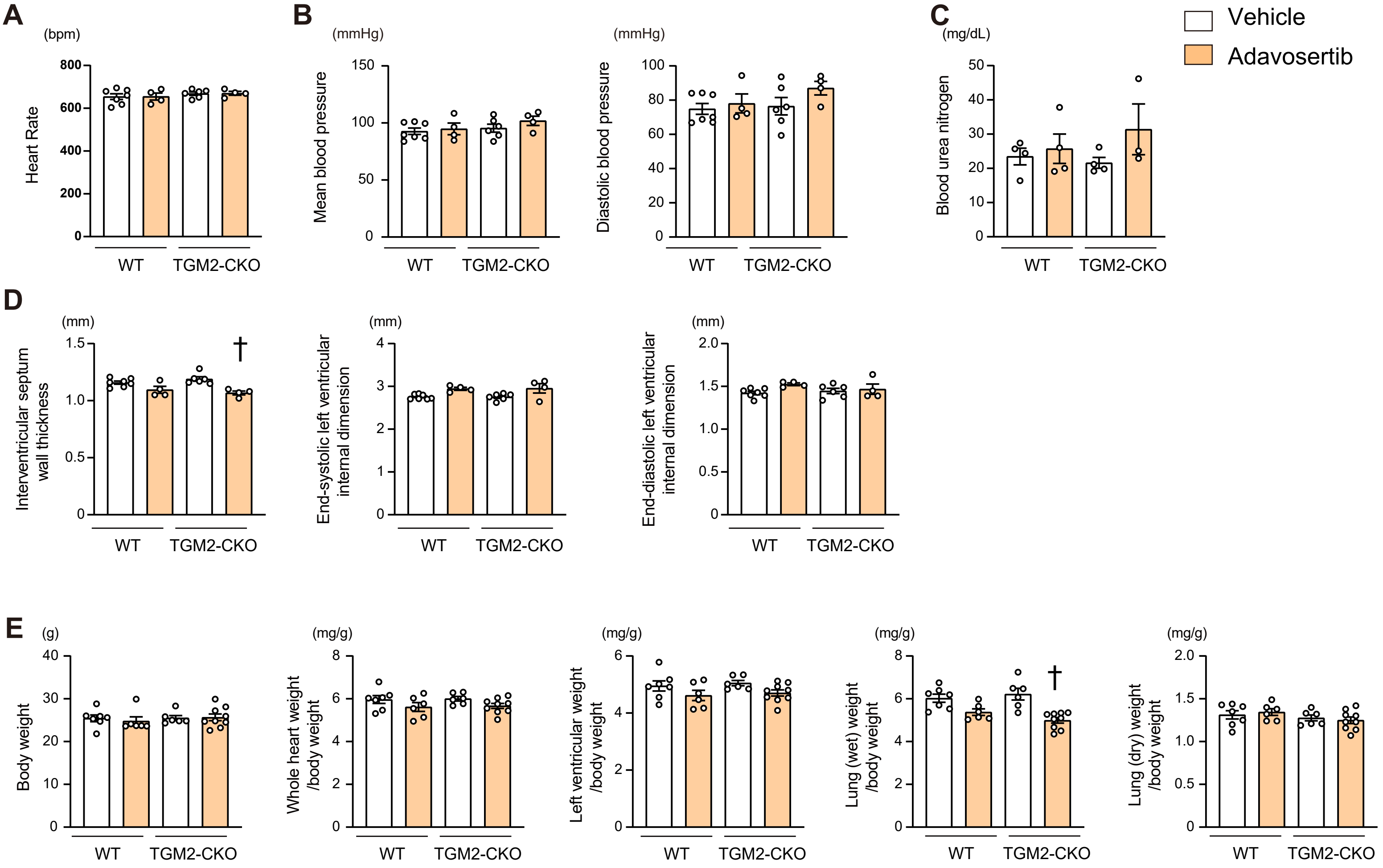


**Supplementary Figure 9. Effect of WEE1 inhibition in SAUNA-exposed TGM2-CKO mice.**
After initiation of the SAUNA model, the WEE1 inhibitor adavosertib (50 mg/kg/day) or vehicle was administered orally once daily for 4 weeks in WT and TGM2-CKO mice. **A**, Heart rate. **B**, Mean and diastolic blood pressure. **C**, Blood urea nitrogen. **D**, Echocardiographic parameters. **E**, Physiological parameters. All data are presented as mean ± SEM. †P < 0.05 versus vehicle-treated TGM2-CKO mice by the one-way ANOVA with Tukey post-hoc analysis.

**Supplementary Table 1. Baseline characteristics stratified by 5-HT levels in patients with HFpEF.**

| | All subjects (n=240) | High 5-HT (> 90.5 ng/mL, n=120) | Low 5-HT (≤ 90.5 ng/mL, n=120) | P value |
|---|---|---|---|---|
| Age, years | 69.1 ± 15.9 | 66.1 ± 17.7 | 72.0 ± 13.3 | 0.004 |
| Male sex, n (%) | 125 (52.1) | 64 (53.3) | 61 (50.8) | 0.698 |
| Body mass index, $kg/m^2$ | 24.0 ± 4.6 | 24.1 ± 4.2 | 23.9 ± 4.9 | 0.737 |
| Systolic blood pressure, mmHg | 126.6 ± 21.2 | 126.2 ± 18.8 | 127.0 ± 23.4 | 0.759 |
| Diastolic blood pressure, mmHg | 70.3 ± 13.5 | 71.0 ± 13.3 | 69.6 ± 13.8 | 0.412 |
| Heart rate, beats/min | 70.1 ± 13.0 | 70.0 ± 12.1 | 70.1 ± 13.9 | 0.957 |
| NYHA functional class III/IV, n (%) | 59 (24.6) | 24 (20.0) | 35(29.2) | 0.099 |
| **Comorbidities** | | | | |
| Hypertension, n (%) | 155 (64.6) | 70 (58.3) | 85 (70.8) | 0.043 |
| Diabetes mellitus, n (%) | 68 (28.3) | 36 (30.0) | 32 (26.7) | 0.567 |
| Dyslipidemia, n (%) | 125 (52.1) | 69 (57.5) | 56 (46.7) | 0.093 |
| Chronic kidney disease, n (%) | 112 (46.7) | 52 (43.3) | 60 (50.0) | 0.301 |
| Atrial fibrillation, n (%) | 100 (41.7) | 46 (38.3) | 54 (45.0) | 0.295 |
| Anemia, n (%) | 93 (38.8) | 36 (30.0) | 57 (47.5) | 0.005 |
| **Medications** | | | | |
| Sodium-glucose transporter 2 inhibitor, n (%) | 15 (6.2) | 9 (7.5) | 6 (5.0) | 0.424 |
| Mineralocorticoid receptor antagonists, n (%) | 38 (15.8) | 20 (16.7) | 18 (15.0) | 0.724 |
| Renin-angiotensin system inhibitors, n (%) | 107 (44.6) | 55 (45.8) | 52 (43.3) | 0.697 |
| Beta blockers, n (%) | 102 (42.5) | 46 (38.3) | 56 (46.7) | 0.192 |
| Diuretics, n (%) | 96 (40.0) | 44 (36.7) | 52 (43.3) | 0.292 |
| Anticoagulants, n (%) | 111 (46.2) | 49 (40.8) | 62 (51.7) | 0.092 |
| Antiplatelets, n (%) | 80 (33.3) | 39 (32.5) | 41 (34.2) | 0.784 |
| **Laboratory data** | | | | |
| B-type natriuretic peptide, pg/mL | 78.2 (35.8-173.7) | 58.6 (26.0-134.2) | 96.0 (46.2-218.7) | <0.001 |
| Hemoglobin, g/dL | 12.4 ± 1.8 | 12.7 ± 1.8 | 12.2 ± 1.8 | 0.04 |
| Platelet, $\times 10^3/\mu L$ | 214.1 ± 84.6 | 214.9 ± 66.8 | 213.2 ± 99.5 | 0.88 |
| eGFR, $mL/min/1.73m^2$ | 60.6 ± 22.1 | 62.6 ± 23.3 | 58.7 ± 20.7 | 0.173 |
| Hemoglobin A1c, % | 5.9 (5.6-6.4) | 5.9 (5.6-6.6) | 5.8 (5.5-6.2) | 0.195 |
| C-reactive protein, mg/dL | 0.48 (0.13-1.07) | 0.46 (0.13-0.97) | 0.51 (0.14-1.24) | 0.383 |
| **Echocardiographic data** | | | | |
| Left ventricular ejection fraction, % | 62.3 ± 5.6 | 62.46 ± 5.18 | 62.17 ± 5.94 | 0.694 |
| Left ventricular mass index, $g/m^2$ | 124.9 ± 37.6 | 120.15 ± 33.24 | 129.63 ± 41.17 | 0.051 |

| | | | | |
|---|---|---|---|---|
| E/e' | 10.3 (7.6-13.9) | 9.98 (7.28-13.3) | 10.7 (8.35-14.6) | 0.047 |
| Tricuspid regurgitation pressure gradient, mmHg | 20.0 (15.0-25.0) | 19.5 (14.0-23.0) | 21.0 (15.0-26.0) | 0.065 |

NYHA, New York Heart Association; eGFR, estimated glomerular filtration rate; E/e', the ratio of early transmitral flow velocity to early diastolic mitral annular velocity. Data are presented as mean ± SD or median (interquartile range). Comparisons between two groups were performed using the unpaired Student's t-test or the Mann–Whitney U test.

**Supplementary Table 2. Correlation between 5-HT levels and other variables.**

| | R | P value |
|---|---|---|
| Age, years | -0.172 | 0.008 |
| Body mass index, $kg/m^2$ | 0.101 | 0.120 |
| B-type natriuretic peptide, pg/mL | -0.225 | <0.001 |
| Hemoglobin, g/dL | 0.127 | 0.050 |
| Platelet, $\times 10^3/\mu L$ | 0.078 | 0.228 |
| eGFR, $mL/min/1.73m^2$ | 0.063 | 0.329 |
| Hemoglobin A1c, % | 0.110 | 0.090 |
| C-reactive protein, mg/dL | -0.068 | 0.297 |
| Left ventricular ejection fraction, % | 0.060 | 0.353 |
| Left ventricular mass index, $g/m^2$ | -0.087 | 0.181 |
| E/e' | -0.071 | 0.273 |
| Tricuspid regurgitation pressure gradient, mmHg | -0.140 | 0.030 |

eGFR, estimated glomerular filtration rate; E/e', the ratio of early transmitral flow velocity to early diastolic mitral annular velocity. Correlations were analyzed by the Spearman correlation test.

**Supplementary Table 3. Univariable and multivariable Cox proportional hazards analyses for cardiac events, including cardiac death and worsening heart failure, in patients with HFpEF.**

| | Hazard ratio | 95% confidence interval | P value |
|---|---|---|---|
| **5-HT as a categorical variable** | | | |
| Low 5-HT (vs. high 5-HT), unadjusted | 3.704 | 1.675 - 8.197 | <0.001 |
| Low 5-HT (vs. high 5-HT), adjusted | 3.247 | 1.441 - 7.353 | 0.005 |
| **5-HT as a continuous variable** | | | |
| 5-HT per 1 ng/mL decrease, unadjusted | 1.013 | 1.005 - 1.020 | <0.001 |
| 5-HT per 1 ng/mL decrease, adjusted | 1.012 | 1.004 - 1.020 | 0.006 |

P values were derived from Cox proportional hazards regression models. Multivariable models were adjusted for age, body mass index, chronic kidney disease, diuretic use, hemoglobin, and B-type natriuretic peptide.

**Supplementary Table 4. Primer sequences used for RT-qPCR.**

| | Gene | | | Sequence | |
|---|---|---|---|---|---|
| Mouse | *Nppa* | Forward | 5’- | TCGTCTTGGCCTTTTGGCT | -3’ |
| | | Reverse | 5’- | TCCAGGTGGTCTAGCAGGTTCT | -3’ |
| | *Myh7* | Forward | 5’- | ATGTGCCGGACCTTGGAAG | -3’ |
| | | Reverse | 5’- | CCTCGGGTTAGCTGAGAGATCA | -3’ |
| | *Col3a1* | Forward | 5’- | CCCGGGTGCTCCTGGACAGA | -3’ |
| | | Reverse | 5’- | CACCCTGAGGACCAGGCGGA | -3’ |
| | *Cdk1* | Forward | 5’- | CTCCTGGGCAGTTCATGGAT | -3’ |
| | | Reverse | 5’- | CCACAGCGTCACTACCTCG | -3’ |
| | *Ccna2* | Forward | 5’- | GGCTGCACCAACAGTAAATCA | -3’ |
| | | Reverse | 5’- | TCAGTTCTCCCAAAAACATTGC | -3’ |
| | *18S rRNA* | Forward | 5’- | GTAACCCGTTGAACCCCATT | -3’ |
| | | Reverse | 5’- | CCATCCAATCGGTAGTAGCG | -3’ |
| Rat | *Cdk1* | Forward | 5’- | CACGGCGACTCAGAGATTGA | -3’ |
| | | Reverse | 5’- | GACCAGCATTTTCGAGAGCA | -3’ |
| | *Ccna2* | Forward | 5’- | TGGATGGTAGTTTTGAATCACCC | -3’ |
| | | Reverse | 5’- | GGCCCGCATACTGTTAGTGA | -3’ |
| | *Nppa* | Forward | 5’- | CTGATGGATTTCAAGAACCTGCT | -3’ |
| | | Reverse | 5’- | GAGAGAGGGAGCTAAGTGCC | -3’ |
| | *Gapdh* | Forward | 5’- | AGTGCCAGCCTCGTCTCATA | -3’ |
| | | Reverse | 5’- | GGTAACCAGGCGTCCGATAC | -3’ |
